# Confinement in fibrous environments positions and orients mitotic spindles

**DOI:** 10.1101/2024.04.12.589246

**Authors:** Apurba Sarkar, Aniket Jana, Atharva Agashe, Ji Wang, Rakesh Kapania, Nir S. Gov, Jennifer G. DeLuca, Raja Paul, Amrinder S. Nain

## Abstract

Accurate positioning of the mitotic spindle within the rounded cell body is critical to physiological maintenance. Adherent mitotic cells encounter confinement from neighboring cells or the extracellular matrix (ECM), which can cause rotation of mitotic spindles and, consequently, titling of the metaphase plate (MP). To understand the positioning and orientation of mitotic spindles under confinement by fibers (ECM-confinement), we use flexible ECM-mimicking nanofibers that allow natural rounding of the cell body while confining it to differing levels. Rounded mitotic bodies are anchored in place by actin retraction fibers (RFs) originating from adhesion clusters on the ECM-mimicking fibers. We discover the extent of ECM-confinement patterns RFs in 3D: triangular and band-like at low and high confinement, respectively. A stochastic Monte-Carlo simulation of the centrosome (CS), chromosome (CH), membrane interactions, and 3D arrangement of RFs on the mitotic body recovers MP tilting trends observed experimentally. Our mechanistic analysis reveals that the 3D shape of RFs is the primary driver of the MP rotation. Under high ECM-confinement, the fibers can mechanically pinch the cortex, causing the MP to have localized deformations at contact sites with fibers. Interestingly, high ECM-confinement leads to low and high MP tilts, which mechanistically depend upon the extent of cortical deformation, RF patterning, and MP position. We identify that cortical deformation and RFs work in tandem to limit MP tilt, while asymmetric positioning of MP leads to high tilts. Overall, we provide fundamental insights into how mitosis may proceed in fibrous ECM-confining microenvironments in vivo.

## Introduction

Eukaryotic cell division proceeds with interphase cells undergoing significant shape changes to become rounded cell bodies. The stiff cortex of spherical cells is held in place by two dominant mechanical cues: retraction fibers (RFs) and mechanical confinement^1–5^. Actin-rich RFs connect the cell body with sites of interphase adhesions, thus acting as stabilizing mechanical links for proper positioning of the mitotic spindle^1,6–8^. The level of RF coverage in fibrous environments leads to diverse mitotic outcomes and errors^9^. On the other hand, confinement from the neighboring cells is ‘volumetric’ and can happen without retraction fibers^10,11^. Improper rounding under confinement can lead to mitotic spindle orientation defects, increased mitotic errors, delayed or halted division, and even cell death^4,11–17^. In loose connective tissues and interstitial matrices having sparse organization of fibrillar collagens (spacing of few to tens of μm)^18–24^, it is conceivable that rounded mitotic cells are held in place by RFs while also mechanically confined between the ECM fibers (we define this to be ECM-confinement). Here, we inquire about the combined action of these cues to direct mitotic spindle positioning and orientation.

Numerous strategies have been developed to study the individual contributions of RFs and confinement on mitosis. RFs are part of the cortical mechanosensory complex, which links the cell cortex to the astral microtubules to transmit the effects of external cues to organize the mitotic spindle^6,25^. Our prior work has shown that increased RF coverage (RFC) on the cell cortex limits rounding cell body movement, causing faster division and a higher incidence of multipolar defects^9^. Studies using flat tissue culture plates and fibronectin-coated micropatterns have shown that RFs formed on the coverslips promote spindle alignment parallel to the substrate^1,2^. On the other hand, when sandwiching (confining) cells between two fibronectin-coated coverslips, the spindles are oriented perpendicularly to the substrate due to the influence of RFs generated on both the bottom and top coverslips^7^. Volumetric confinement using an AFM cantilever placed directly on the rounded cell has shown that low confining forces (∼ 5 nN) accelerate mitotic progression, while at larger confining forces (∼100 nN), the integrity of the mitotic spindle and the overall mitotic progression is impaired^4,5^. Maintaining proper orientation of mitotic spindles under confinement requires appropriate tension levels at intercellular junctions in the epithelium plane^26,27^, but increasing the confining forces by using dense gels leads to delayed division^10^, buckling of mitotic spindles^28^, and spindle multipolarity defects leading to cell death^14^.

Here, we introduce a repeatable system of suspended nanofibers mimicking the sparse extracellular environment that is capable of interrogating the combined effects of ECM-confinement and RFs in orienting the mitotic spindle. The suspended and flexible nature of fibers allows the natural rounding of cells undergoing division, confining them to differing levels while mechanically linking the cortex and interphase adhesion sites through RFs. We find that the extent of confinement regulates the 3D patterning of RFs on the cortex, and high confinement causes localized deformations of the cell cortex and, consequently, in the metaphase plate. For a mechanistic interpretation of the outcome due to the combined effects of cortical deformation and RF patterning, we developed a stochastic Monte Carlo *in silico* framework accounting for the centrosome (CS), chromosome (CH), and membrane interactions. We discover that the patterning of RFs primarily governs the metaphase (MP) tilt, and the cortical pinching creates physical constraints on MP positioning. Overall, we provide knowledge of the cooperative and competing effects of ECM-confinement and RF patterning in orienting the mitotic spindle, as would be expected in fibrous in vivo environments.

## Results

### Mechanical confinement in fiber networks enables measurement of mitotic forces and induces tilt of the metaphase plate

Mitotic cells exert outward forces as they round up against volumetric confinement. In our previous work, we showed that cells attached to two fibers (12 μm apart) underwent division by balling up while being held by RFs originating from four adhesion sites on fibers (**Fig. 1a(i)**, *Supplementary Movie S1*) ^9^. Often the cells got trapped between the two fibers, causing them to bow outwards (**Fig. 1a(i)**, *Supplementary Movie S2*), which we termed as ECM*-*confinement. Naturally, we inquired if our model system could be used to study the combined effects of the extent of confinement and RF arrangement on orienting the mitotic spindle and metaphase plate (MP). To quantify the extent of confinement, we analyzed the basal-apical positioning of the rounded cell with respect to the fiber plane from the confocal cross-sectional side views (yz, **Fig. 1a(ii)**). We measured two heights: cell height (H) and height of fiber plane (h) from the bottom of the cell, with a value of h/H = 0 indicating a pure unconfined cell positioned on top of fibers (no bowing out of fibers), and h/H = 0.5 showing confinement at the cell mid-plane with the maximum outward bulge of fibers and formation of two cellular lobes. Thus, we categorized cells as confined for h/H values ranging from 0.3 to 0.5 and unconfined for h/H ratios ranging from 0 to 0.3, **Fig. 1a(iii)**. Mechanical confinement resulted in noticeable cortical deformations at sites of contact with fibers (yellow arrowheads in **Fig. 1a**), which was associated with increased cell height (**Fig. 1a(iv)**, *Supplementary Fig. S1a*) and aspect ratio (*Supplementary Fig. S1b*).

**Figure 1:**
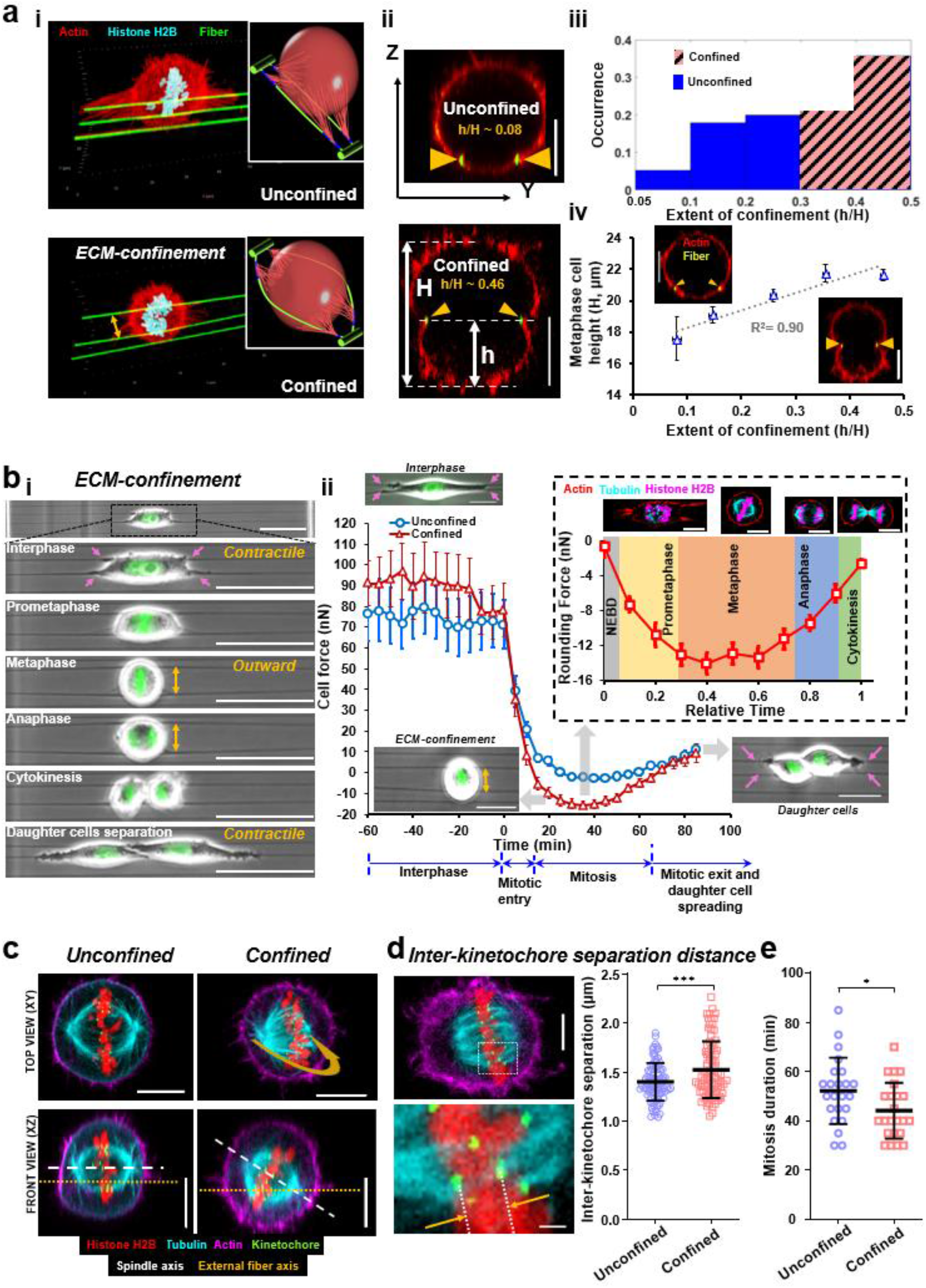
Mechanical confinement of mitotic cells in fibrous environments, a) i) Representative 3D isometric views of single mitotic cells on top of fibers (unconfined) or trapped between 2 fibers (confined), actin (cell cortex), histone H2B (chromosomes) and the fibers (fibronectin coating) are indicated in red, cyan and green respectively. Inset cartoons highlight the outward bowing of fibers during ECM confinement. ii) Representative side-views (YZ cross-section) of unconfined and confined cells with the fibers as green dots (yellow arrowheads), H represents the cell height taken at metaphase, h is the height from the fiber plane to the bottom of the cell, scale bars are 10 μm, iii) Histogram showing relative occurrence of the different levels of confinement (quantified by h/H, n= 95), iv) Metaphase cell height increases with confinement (h/H) (R^2^ = 0.9, p = 0.0132). Representative cross-sectional images with very low h/H (∼0.1) and high h/H (∼0.5). Scale bars are 10 μm. b) i) Timelapse images of a representative confined cell undergoing cell division, ii) Force profiles of confined and unconfined cells transitioning from interphase to mitosis, with inset images showing a single cell over different stage. By convention, mitosis forces are shown as negative to represent outward deflection of fibers. The dashed rectangle box shows the average force profiles for confined cells rounding during mitosis which is normalized for time taken from NEBD to cytokinesis (n=23) along with representative top views of cells undergoing various stages of division. c) Representative top (xy) and front (xz) views of confined and unconfined cells demonstrating the 3D tilt of the metaphase plate (MP). Yellow arrow represents the MP rotation. Cells are stained with actin cortex (magenta), microtubules (cyan), kinetochores (green) and chromosomes (red). Scale bars represent 10 μm. d) Representative images showing the measurement of inter-kinetochore separation and the quantitation showing an increase in the average interkinetochore distance with confinement. Scale bars represent 5 μm. e) Confinement causes a drop in the mitosis duration, n=47.

The bending of fibers allowed us to estimate the forces during interphase, mitosis, and post-cell division (**Fig. 1b (i, ii)**, *Supplementary Movie S3*) using Nanonet Force Microscopy (NFM, *Appendix I in Supplementary Materials*)^29–31^. During interphase, cells are in their natural contractile state and thus deflect the fibers inward (pink arrows, **Fig. 1b (i, ii)**. Contractile forces are, by convention, considered to be positive. Following mitotic entry in confined conditions, cells progressively round up by forcing the fibers outwards (yellow arrows, **Fig. 1 b (i, ii)**). Consistent with literature-reported data, we observed a steady increase in the magnitude of the rounding forces from nuclear envelope breakdown, with peak mitotic forces of ∼12-14 nN achieved during metaphase ^5,32^. In contrast, in the case of unconfined cells, we did not observe appreciable expansive forces (outward deflection of fibers). Since RFs anchored the cells, our force measurement platform allowed us to estimate the tension in RFs. We estimated the tension in RFs (∼214 pN) by assuming RFs from each adhesion site to be analogous to springs-in-parallel (average number of RFs ∼10 per focal adhesion cluster (FAC), *Supplementary Fig. S2*). Following the mitotic exit, we observed that daughter cells started spreading and re-applying contractile forces (**Fig. 1b (ii)**).

Next, we investigated if the ECM-confinement affected the mitotic spindle’s and MP’s organization. We found that ECM-confinement caused the mitotic spindles to rotate inside the cell body with a large MP tilt visualized from the xz-front views of confocal z-stack images (**Fig. 1c**). Additionally, ECM-confinement correlated with an increase in the average inter-kinetochore separation distance during metaphase (**Fig. 1d**) and significantly faster mitosis (**Fig. 1e**). The observation that increased inter-kinetochore separation distance corresponds to faster mitotic durations matches our previous observations^9^. Cytokinesis events captured from live cell imaging revealed substantial reorientation of the division axis in cells under ECM-confinement (*Supplementary Fig. S3, Supplementary Movie S4*).

Overall, we quantified the mitotic forces as cells balled up under ECM-confinement. We found that ECM confinement caused tilting of the MP and reduced mitotic durations.

### The patterning of retraction fibers governs MP tilting

We wanted to interrogate the factors responsible for the tilt of the MP under ECM-confinement. We identified two key parameters: i) the extent of mechanical confinement and ii) the 3D organization of the RFs. Using confocal microscopy, we generated 3D z-stacks of fluorescently labeled fibers, cell cortex, and the MP (**Fig. 2a (i, ii)**), which allowed us to correlate the increase in MP tilt from the xz-front views with the extent of confinement from its yz-side views (R^2^=0.96, **Fig. 2a (iii)**).

**Figure 2:**
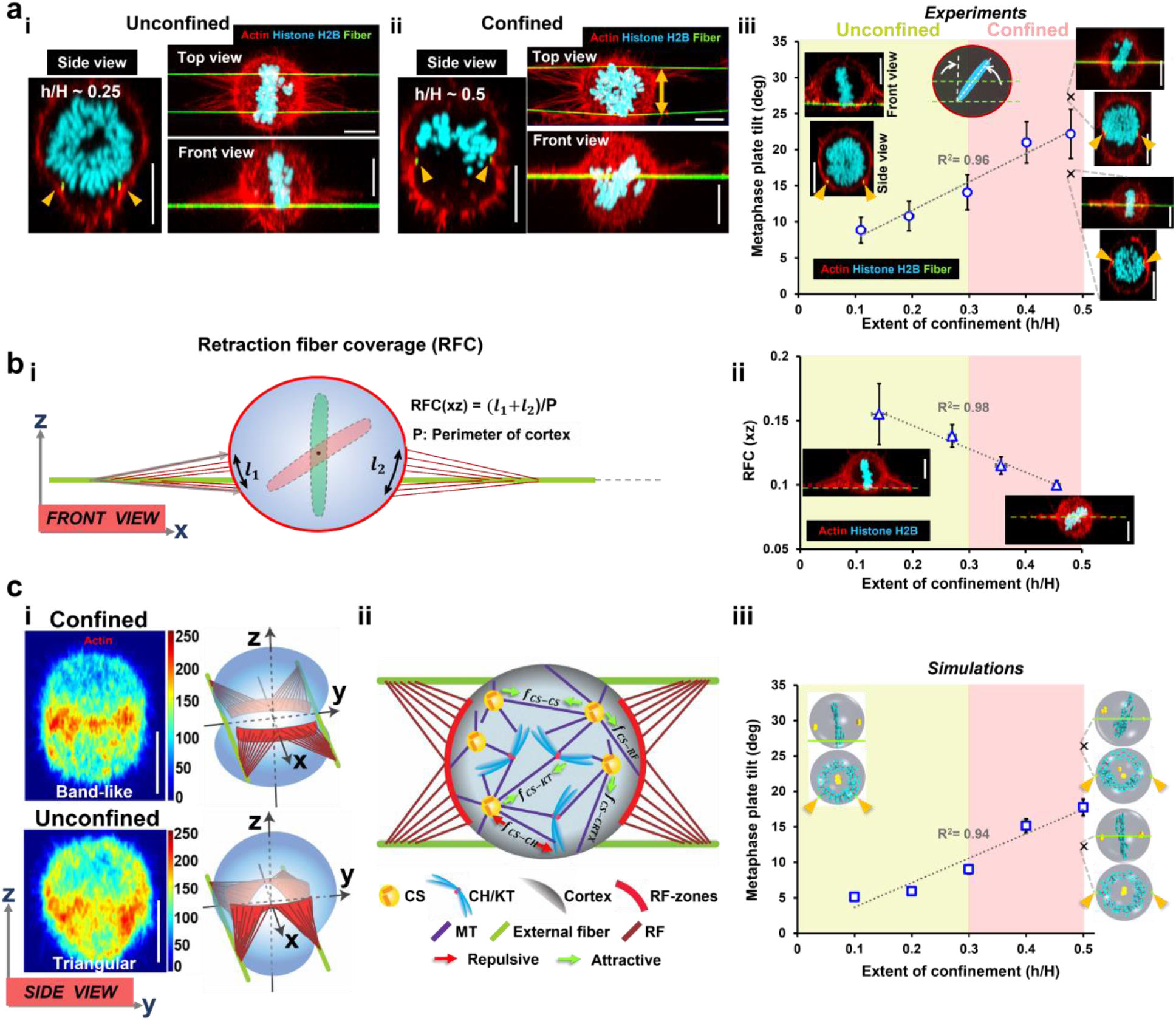
Computational modeling recapitulate experimentally observed MP tilt with increasing confinement, **a) i, ii)** Representative unconfined and confined cells with top (xy), side (yz) and front (xz) views shown, actin, histone H2B and fibronectin are shown in red, cyan and green to identify the cell cortex, metaphase plate and the fibers respectively, scale bars are 10 μm, **iii)** Metaphase plate tilt as a function of the extent of confinement (h/H), inset images shows front and side views of unconfined (h/H∼ 0.1) and confined (h/H ∼0.5) cells. The angle is measured from a normal drawn to the fiber axis (green dashed lines shown for confined and unconfined cases). Scale bars are 10 μm, **b i)** Schematic showing the quantification of the retraction fiber coverage (RFC), **ii)** Retraction fiber coverage decreases with h/H, inset images showing RF organization in representative front views of confined and unconfined cells, scale bars are 10 μm, **c i)** Intensity heatmap demonstrating the average projection of yz-side views in actin stained cells, scale bars are 10 μm, Band-like arrangement emerges for RF organization in confined cells, while unconfined cells have triangular forms of RF regions that extends downwards from the mid-cortical level, corresponding adopted RF regions for the computational simulations, **ii)** Computational model of the mitotic cell between the 2 external fibers (marked in green), CSs attract each other (*f*_*CS*−*CS*_) or KTs (*f*_*CS*−*KT*_) and are attracted to the cortex region devoid of RFs (*f*_*CS*−*CRTX*_) or coupled to RFs (*f*_*CS*−*RF*_), CSs repel chromosome arms (*f*_*CS*−*CH*_), all relevant components of the mitotic machinery are marked and indexed below, *f*_*CS*−*RF*_ attraction is considered several times stronger than *f*_*CS*−*CRTX*_, **iii)** computational data for MP tilt as a function of h/H, representative snapshots showing the front and side views of spherical unconfined and confined cells. Note that crosses in a(iii) and c(iii) show the low and high MP tilt angles observed at high-confinement along with representative images. The average of these two angles are plotted in the main figure and the origin of two configurations is explained in Figure 3.

Next, we analyzed the spatial patterning of RFs under different levels of ECM-confinement. From the xy-top views, we observed that both confined and unconfined cells were held in space by four sets of RFs attaching the cell cortex region between the external fibers (top view in **Fig. 2a (i, ii)**). Such an arrangement originates from the adhesion geometry of these cells during interphase in agreement with our previous study^9^. We observed that the patterning of RFs extended from the fiber plane to the cells’ mid-cortical (equatorial) level (*Supplementary Fig. S4*). To quantify ECM-confinement-driven RF coverage, we measured the proportion of the cortical perimeter (xz-view) covered by retraction fibers (**Fig. 2b (i)**). We found RF coverage decreased with increasing levels of confinement (h/H, **Fig. 2b (ii)**), signifying a band-like and triangle-like pattern of RFs at high and low confinement, respectively (**Fig. 2b (ii**, inset) and **Fig. 2c (i)**).

Next, we sought to mechanistically explain the underlying mechanisms causing the variation in the MP tilt as a function of the extent of confinement (h/H) and the retraction fiber patterns. We developed a computational model to account for centrosomes (CSs), chromosomes (CHs), kinetochores (KTs), and cell cortex in a rounded mitotic cell. The model is built upon our earlier framework^9,33^, with augmentation to include spatial organization of experimentally observed RF patterns during ECM-confinement. We simulated a pairwise interaction model where the model entities (CSs, CHs, KTs) interacted among themselves and with cortex through forces that vary with the separation distance ^33–36^. However, the forces between CS and KT are assumed to be distance-independent ^33,37^. Thus, our model includes five fundamental forces: CS attraction to each other, CS attraction to the KTs, CS repulsion from the CH arms, and short-range attraction among CSs to the RF regions and the remaining cortex (CRTX) (**Fig. 2c (ii)** and details of the model are included in *Appendix II* in *Supplementary Materials*). These forces arise from microtubule dynamics and molecular motors interacting with various model components^36^. The CS-CS, CS-CH, and CS-KT forces were maintained at a base value corresponding to a bipolar majority^33^. The interaction forces between CS and RFs transmitted through the astral microtubules (MTs) were set several times stronger than the forces from the remaining CRTX (see *Table II* in *Supplementary Materials* for model parameters). This is consistent with our earlier observation that the RFs interact with the CSs ∼4 to 10 times stronger than the remaining CRTX. Such consideration is crucial for achieving the observed propensity of spindle statistics in cells undergoing mitosis in various fibrous environments^9^. Coupling the five force interactions in a stochastic energy minimization framework, we performed Monte Carlo simulations to achieve stable mechanical equilibria of the spindle.

Next, we investigated the mechanism behind the MP tilt under ECM-confinement computationally. Our experiments showed that the extent of confinement correlated with changes in cell height and patterning of RFs in band-like and triangular shapes at high and low confinement, respectively. We recreated the RF patterns in our model (**Fig. 2c (i)**; see simulated RF pattern in *Table I, Case 5* in *Supplementary Materials*). We determined that the contribution of cell height on MP tilting was insignificant (*Supplementary Fig. S5*). Our computational model successfully reproduced the observed trend of increased MP tilt with increasing levels of confinement (**Fig. 2c (iii)**). The model suggests that the smaller MP tilts at low ECM-confinement results from the increased RFC area (**Fig. 2b (ii)**) due to the triangle-like patterning of the RFs. The increased RFC causes a substantial pull on the CSs from the RF regions, yielding lower MP tilt. Among numerous simulated RF patterns of distinct shapes and locations on the spherical cortex (*Table I* in *Supplementary Materials*), the only configurations that matched the experimentally observed trend of MP tilt coincided with triangular RF patterns at low confinement. For cells under high ECM-confinement (h/H ∼0.5), we observed low and high MP tilt angles (shown by crosses in **Fig. 2a(iii)** and **Fig. 2c(iii)**). Such distinct variations in the MP tilt angles were unexpected, and we expand upon their origin in the next section.

Therefore, by combining experiments and mechanistic computational modeling, we demonstrate that patterning of RFs (triangle shape in unconfined and band-like in confined) is the dominant factor controlling the MP tilting.

### The interplay of RFs and cortical pinching regulates the extent of MP positioning and tilt under high ECM-confinement

Next, we inquired about the origin of low and high MP tilt angles at high ECM-confinement (h/H ∼0.5). We found that high ECM-confinement by the parallel fibers could pinch the cell, forming two symmetric cellular lobes due to local deformation in the cortex. Thus, we interrogated the independent contributions of cortical pinching and RFs in tilting the MP under high ECM-confinement. A close examination of the confocal front views revealed the MP plate to be symmetrically (similar proportion in both cellular lobes) and asymmetrically (major proportion in one cellular lobe) positioned (**Fig. 3a (i, ii)**). We developed a new metric *h*_*f*_ (**Fig. 3a (iii)**), denoting the distance between the MP centroid and the fiber plane to describe the proportion of MP in cellular lobes: a value closer to zero signifying equal proportion in both lobes. From our experiments and simulations, we observed a higher occurrence of cells with low *h*_*f*_ (< 1 μm) that were associated with low MP tilt angles (**Fig. 3a (iv)** and *Supplementary Fig. S6*). However, for *h*_*f*_ values greater than 1 μm (MP positioned asymmetrically), we observed a sharp increase in the MP tilt angle (**Fig. 3a (iii, iv**)). Inspection of confocal images of *h*_*f*_ >1 μm cases revealed one of the spindle poles was positioned far from the RF regions, while the other pole remained close to the band-like RF regions on the opposite cell side (see Figure 1c (confined case front view) and *Supplementary Fig. S7*).

**Figure 3:**
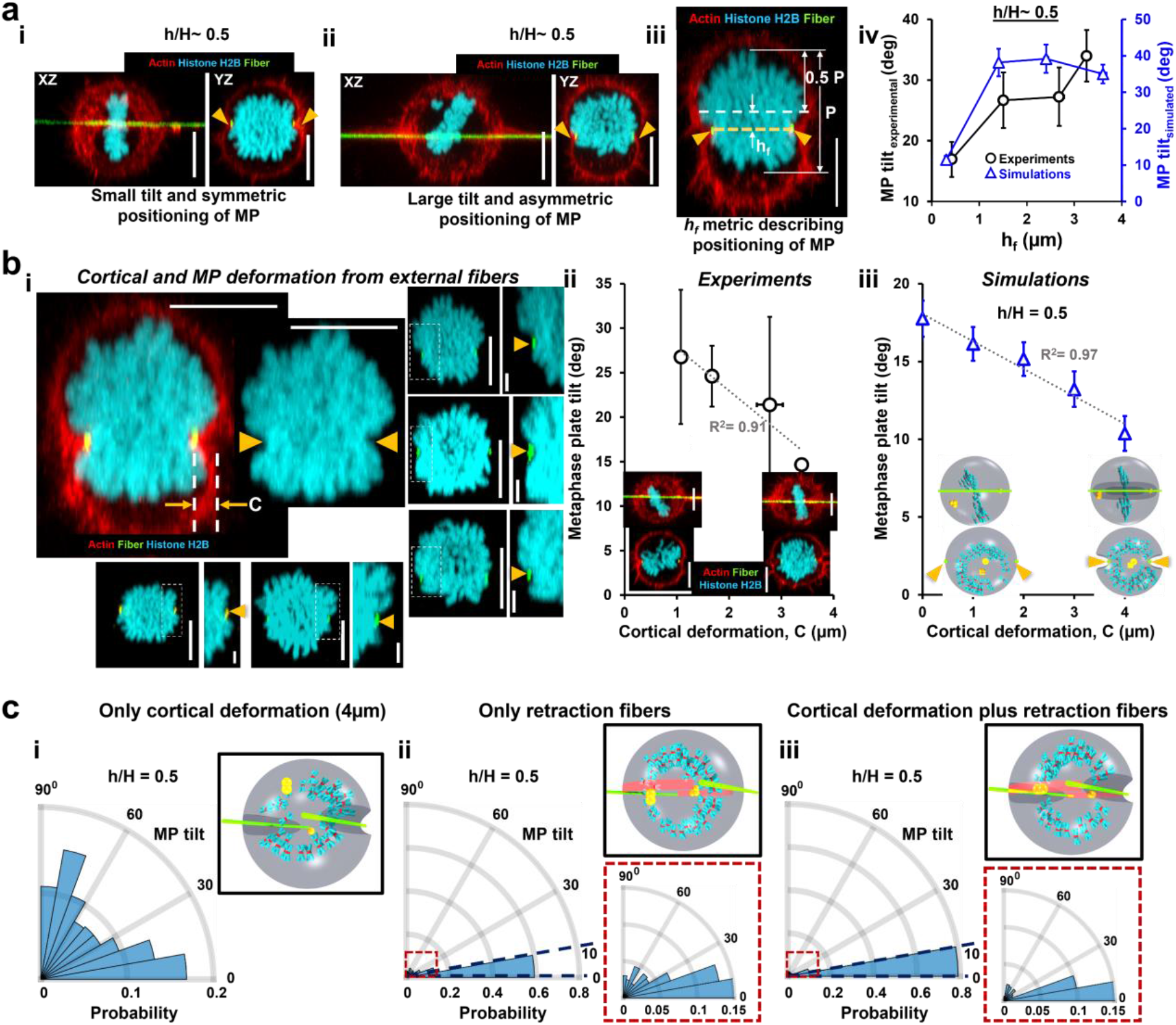
Relative influence of pinching and RF distribution on the MP tilt angle in confined cells. **a i**,**ii)** Front (xz) and side (yz) views of representative confined cells with similar confinement level (h/H ∼ 0.5) but with small and large MP tilt respectively, **iii)** Representative side view of a confined cell showing the *h*_*f*_ metric, which denotes the distance between the center of the metaphase plate and the plane of the fibers, ‘P’ denotes the length of MP, and 0.5 P defines the MP mid-plane, **iv**) MP tilt angle as an increasing function of *h*_*f*_, as observed in the experiments and also confirmed by computational modeling, **b i)** Side-view of a representative confined cell demonstrating the cortical deformations (*C*) induced by the external fibers. *C* is measured as the horizontal distance from the fiber position to the line tangent to the cortical surface of the cell. Inset demonstrates that cortical deformations can further lead to deformations (denoted by yellow triangles) within the metaphase plate. Five more representative cases showing deformations in the cortex and MP, with scale bars of 2 μm in magnified views of boxed regions. **ii)** Average MP tilt from experimental observations as a function of cortical deformation, C, **iii)** MP tilt as a function of cortical deformation as generated from simulations, **c)** Polar histograms of the MP tilt angles generated from computational simulations in the presence of **i)** only cortical deformation (with C = 4 μm) **ii)** only retraction fibers and **iii)** presence of both cortical deformations (C = 4 μm) and retraction fibers, Insets in ii and iii demonstrate a magnified view of the polar histograms, CSs are in yellow, CHs are blue, KTs are in red, and two green lines denote external fibers; RF attachment areas are represented as two light red bands on the cell surface. All scale bars represent 10 μm, unless noted.

Next, our confocal images revealed large deformations at sites of pinching of the cortex by external fibers (**Fig. 3b (i)**). We found that these deformations led to sharp changes in MP at sites of pinching from the external fibers (**Fig. 3b(i)** more representative examples), which suggests a physical limitation in the translation of MP. We measured the extent of deformation from outside the cell cortex (*C* in **Fig. 3b (i)**) and found higher cortical deformations correspond to lower tilt angles (**Fig. 3b (ii)**). On the other hand, unconfined cells having negligible cortical pinching showed minimal MP deformation (*Supplementary Fig. S8)*. Next, we investigated how cortical deformation influenced the positioning of the MP and, consequently, its tilt. The cortical deformation imposes geometrical constraints on the CSs/CHs movement. Our simulation data revealed that increasing values of cortical deformation correlated with *h*_*f*_ *<* 1 μ*m*, signifying symmetric distribution of MP in both cellular lobes (*Supplementary Fig. S9*). Our experimental findings partially support these observations, likely due to cell heterogeneity and inadequate cases of large cortical deformations (*Supplementary Fig. S10*). Similar to findings from our experiments, our simulations showed reduced MP tilt with increased cortical deformation (**Fig. 3b (iii)**).

Next, we used our computational framework to examine the cooperative ability of cortical deformation and RFs to regulate the MP tilting under high ECM-confinement. First, we examined the effect of RFs by themselves, followed by the combined effect of RFs and cortical deformation. In the absence of cortical deformation, we note that at *h*_*f*_ ≤ 1 μm, the reduced MP tilt is due to stronger CS-RF attraction that counters the CS-CRTX attractive forces from the remaining cortex, thus aligning the spindle parallel to the fiber plane. At *h*_*f*_ > 1 μm, signifying the positioning of MP in one of the cellular lobes, the observed high MP tilt is due to one CS pole positioned far from RF regions driven by the CS-CRTX attraction, while the other pole remains close to RFs on the opposite cell side influenced by the CS-RF attractive forces. Next, we examined the role of cortical deformations without and with the presence of RFs. We reasoned that high cortical deformation near the fiber plane (*h*_*f*_ ⩽ 1 μm) physically limits the translation of the MP in one of the cellular lobes, thus significantly reducing the tilting as well. We discovered that without RFs (i.e., with uniform distribution of cortical cues for CS-CRTX interactions), pinching was inadequate for aligning the spindle with the fiber plane, resulting in a broad distribution of MP tilt angles (**Fig. 3c (i)**). On the contrary, without pinching, RFs were sufficient to align most spindles with the fiber planes (**Fig. 3c (ii)**), as described above. However, the absence of cortical deformation cannot rule out the translation of MP (*h*_*f*_ > 1 μm), which resulted in a small fraction of cases displaying tilted MPs, as discussed above (inset in **Fig. 3c (ii)**). Combining the effects of cortical deformation and RFs in the simulations resulted in an MP tilt similar to that of RFs alone (**Fig. 3c (iii)**). However, both cues working in tandem result in a rise in the proportion of cells with low MP tilts (≤ 10^0^). Our simulations at different values of cortical deformation further elucidate the cooperativity of RFs and cortical deformations under two conditions (*h*_*f*_ *<* 1 μm and *h*_*f*_ > 1 μm, *Supplementary Fig. S9*). At *h*_*f*_ *<* 1 μm, we observed a decrease in MP tilt angle with increasing cortical deformation, while at *h*_*f*_ > 1 μm, we observed high tilt angles independent of cortical deformations.

Overall, we show that the MP tilt is sensitive to the positioning of MP relative to the fiber plane. MP centroid away from the fiber plane exhibits high MP tilt due to the reduced influence of RFs. MP centroid close to the fiber plane predominantly exhibits low MP tilts due to the influence of RFs as the primary mechanical cues for spindle alignment. Cortical deformation, which physically limits the movement of MP, further enhances the proportion of cells displaying low MP tilt angles.

## Discussions

We report how mechanical ECM-confinement impacts the positioning and orientation of mitotic spindles. We generate a repeatable system of cells undergoing division using aligned nanofiber networks of inter-fiber spacing 12 μm, allowing the mitotic cells to confine between the external fibers at varying levels. In the present model, we found a significant proportion (∼55%) of cells entrapped between the fibers while rounding up for mitosis. The fibers are suspended and flexible, which allows the natural rounding of cells. Thus, our approach establishes a method to study the mitotic outcomes under confinement as expected in sparse fibrous in vivo environments of loose connective and interstitial tissue with interfiber spacing ranging from a few to 20 μm^18–24^. While rounding up, cells deform the fiber networks while being held in position by RFs that connect the cell cortex to the adhesion sites. We find that the extent of confinement patterns RFs on the cortex and the deformed ECM fibers apply a restorative compression to cause a local deformation in the cell cortex and, consequently, in the MP. By combining quantitative microscopy and computational modeling, we mechanistically interrogate the combined effects of confinement-driven RF patterning and cortical deformation in regulating the MP positioning, orientation, and mitotic duration.

While mechanical confinement and cell-ECM/cell-cell connections have been individually shown to influence mitotic spindle orientation, their combined effects remain poorly understood. Disruption of RF-mediated connections in unconfined cells through pharmacological treatment or laser ablation results in mitotic spindle misalignment^1,6,7^. Using a vertical microcantilever array, Sorce et al. demonstrated that confined cells with insufficient rounding force (impaired actin cytoskeleton) can undergo large spindle rotations (∼70°)^32^. *In vivo*, confined mitotic cells behave similarly, with an almost orthogonal reorientation of the mitotic spindle upon loss of actin cortical tension^38^. At the cellular level, two major forces dominate: i) the outward rounding forces at the start of cell division caused by a multifold increase in the cortical tension^39^ and the intracellular hydrostatic pressure^5^ and ii) the pulling forces exerted on the rounded cell body, either from the neighboring ECM (actin-rich RFs)^1,2,40,41^ or neighboring cells (cell-cell cadherin based connections) in a tissue^41–43^. Mitotic rounding forces counteract the externally imposed mechanical confinement from surrounding ECM or cells and have been previously measured in vitro via AFM cantilevers and vertical cantilever arrays^4,5,44^. Contractile forces existing within RFs have been characterized via optical tweezers. Our deformable nanonets allow us to quantify the mitotic rounding forces while indirectly measuring the forces within each RF. A caveat in our force measurement is that forces are calculated from the top view; thus, we cannot relate the force values with the extent of confinement. However, mitotic force calculation of 12-14 nN is similar to the ∼15 nN peak force for HeLa cells reported using the vertical cantilever assay^32^. Furthermore, our estimate of RF tension (∼214 pN) from cells not completely rounded agrees with previously reported values using optical tweezers (250 pN)^1^.

We observed MP tilting and positioning depended upon the extent of confinement. To gain a mechanistic understanding of these outcomes, we utilized our computational model, encompassing various MT-dependent forces regulated by internal and external factors involved in mitotic spindle organization. Numerous studies have shown that cortically localized protein complex composed of Gαi, LGN, and NuMA facilitate the recruitment of the MT minus-end-directed dynein-dynactin complex to the cell cortex, exerting pulling forces on the astral MTs extending from spindle poles^45,46^. External factors of RFs and confinement can significantly alter spindle orientation by causing a reorganization of the internal molecular components and enriching them in the cortical region attached to RFs^6–8,25,41^ or the intercellular junctions^26,41,42,47^. In addition to Gαi/LGN/NuMA, other cortical mechanosensory proteins, such as integrins, FAK, Src, p130Cas, caveolin-1, and MISP, have been found to enrich near the cortex region connected to RFs, exerting a significantly stronger pull on the CSs via the astral MTs than the rest of the cell cortex^6–9,25^. Our simulations account for experimentally observed cortical cues in RF-covered and RF-free cortical regions through CS-RF and CS-CRTX attractive forces, respectively (see **Fig. 4a**). Notably, in the unconfined cells (low ECM-confinement), positioning the rounded cell body at or above the fiber plane results in a triangular pattern of RFs that cover a large area on the cell cortex spanning from the fiber plane (sites of adhesions) to the equatorial plane of the cell body. Thus, we expect that a larger RF coverage area will attract evolving CS clusters on opposite cell halves towards the corresponding RF regions with enhanced CS-RF attractive forces, which will subdue the effect of the CS-CRTX attraction from the remaining cortex. The larger RF regions, in turn, align the mitotic spindle parallel to the equatorial plane (analogously to the external fiber plane), yielding smaller MP tilts. On the other hand, in confined cells (high ECM-confinement), the band-like arrangement of RFs at or near the equatorial plane with reduced RF coverage results in a relatively weaker influence of the CS-RF forces while enhancing the net CS-CRTX attraction arising from the rest of the cortex. These competing forces acting on the CSs lead to more MPs acquiring a greater tilt angle than the unconfined cells (*Supplementary Fig. S11*). Furthermore, our computational model reveals a surprising role of the triangular shape of RF patterns in yielding smaller MP tilt angles at low confinement. Within a diverse array of simulated RF patterns on the cortex, the only configurations matching the experimentally observed trend of MP tilt are those with triangular shapes. Conversely, deviations from triangular RF patterns, such as rectangular RF spots or compact band-like patterns extending from the fiber plane to the cell equator at low ECM-confinement, lead to MP tilting, which is inconsistent with experiments (*Table I, Case 2-4* in *Supplementary Materials*).

**Figure 4:**
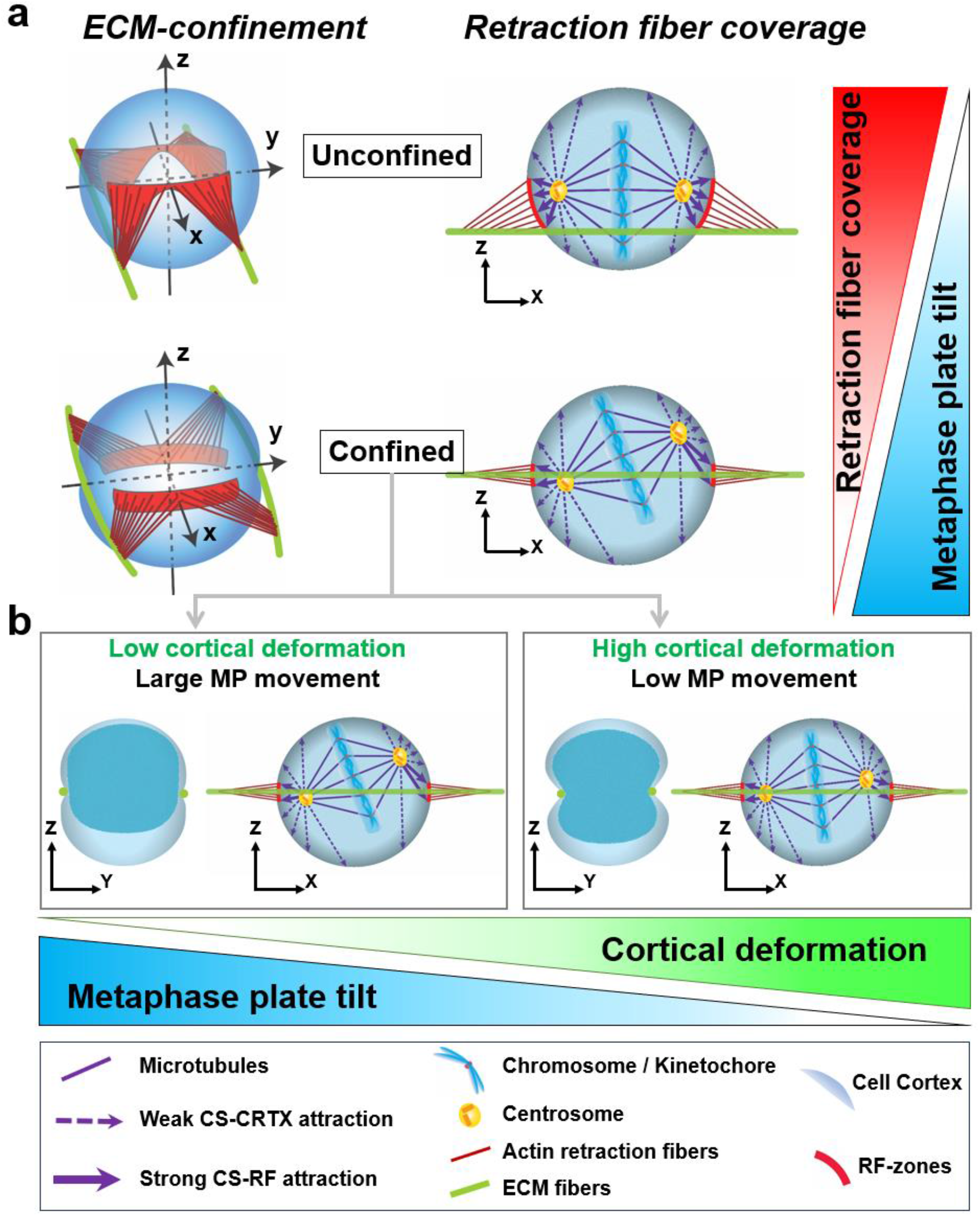
A schematic representation of model outcomes outlining the role of retraction fiber organization and cortical deformation regulating MP orientation. Front (xz) views of the cell show MP tilt and corresponding spindle orientation. (a) The decreased RFCs with band-like RF organization induce increased occurrences of large MP tilts in confined cells compared to unconfined cells having triangular RF organization. (b) The cortical deformation in confined cells imposes physical constraints on MP movement, reducing the overall MP tilt with increased cortical deformation. Low cortical deformation, with no MP deformation, generally allows for both translational and rotational MP movement, resulting in a high MP tilt. Conversely, higher cortical deformation accompanied by localized MP deformation, lowers MP tilt by restricting its translational movement away from the fiber plane and rotation prompted by CS-CRTX attraction.

In addition to the small RF coverage due to band-like RF patterning, cells at high confinement exhibit significant cortical deformation caused by external fibers. We investigated whether such deformation in the cell cortex can subdue the effect of low RFC that contributes to the tilting of the MP. Indeed, our experiments and computational studies show that increased cortical deformation, leading to localized MP deformation, correlates inversely with MP tilt angles. Deformation constrains MPs to be physically limited in movement, which works in tandem with RF cues to outweigh the CS-CRTX attractions from the remaining cortex, causing spindle misalignment (tilted MP) (see **Fig. 4b**). To determine the cooperative ability of cortical deformation and RF cues, we simulated our model under three conditions: cortical deformation only, RF cues only, and a combination of both these cues. The MP tilt angles are random under cortical deformation alone (without RFs), consistent with previous findings in cells cultured on poly-l-lysine (PLL) that cannot establish RFs and, thus, display random orientations of the mitotic spindle^6,7,48^. However, the sole presence of RFs in the model caused more cells to have low MP tilts. Combining both cues (cortical deformation and RFs) increased occurrences of MPs with low tilt. Altogether, our study reveals a clear correlation between the positioning of MPs relative to the fiber plane and the resulting MP tilt angles. MPs located close to the fiber plane predominantly exhibit reduced tilt angles due to the influence of CS-RF attraction and physical limitation in the movement of MP due to pinching from the external fibers. On the other hand, MPs positioned further away from the fiber plane tend to have higher MP tilt angles due to dominant CS-CRTX attractive forces from the remaining cell cortex.

Interestingly, cells under high ECM-confinement correlate with faster mitosis times. This observation is suggestive of the rapid formation of stable KT-MT attachments and faster silencing of spindle assembly checkpoint (SAC) prior to metaphase/anaphase transition. Different studies have identified the presence of unattached kinetochores and tension on kinetochores as potential regulators of SAC activity during mitosis^49–53^. Laser-ablation of the last unattached kinetochore was shown to facilitate faster transition of the cells to anaphase^49^, suggesting that delayed KT-MT attachment can facilitate the SAC to remain activated for a longer period, thereby prolonging mitosis. Compared to confined cells where CSs tend to position on the opposite cell sides near the mid-cortical level under the influence of RFs on the equatorial cell surface, the unconfined cells with RFs distributed over a broader region below the mid-cortical level promote the CSs to place much below the mid-cortical level. Such centrosomal arrangement at low ECM-confinement may delay in attachment of MTs to chromosomes located in the hemisphere lacking the CSs, contributing to the prolonged activation of SAC. Furthermore, recent studies have also suggested the role of KT tension in regulating mitotic timing by correcting attachment errors. Increased tension on kinetochores, resulting from microtubules pulling from opposite spindle poles, induces phosphorylation of certain kinetochore proteins, facilitating the release of incorrectly attached microtubules to achieve bi-orientation, followed by a transition to anaphase ^50–52,54^. Therefore, the faster mitosis, correlating with larger inter-KT distance at high confinement can emerge as a consequence of larger tension on the kinetochores in confined cells. The substantial inter-KT distance at high ECM-confinement might be attributed to the MPs positioned near the fiber plane, aligning the spindle axis with external RF regions on opposite cell surfaces at the mid-cortical level. This linear arrangement potentially facilitates greater axial tension (along the spindle axis) applied to the centrosomes from the RF regions, leading to an enhanced inter-KT distance. In contrast, at low ECM-confinement, centrosomes are connected to RF regions distributed below the mid-cortical level. This specific arrangement reduces effective tension on the centrosomes, lowering the inter-KT distance. However, the current model and experiments have limitations in verifying these possibilities and would invoke future investigations.

Our results on cell division under ECM-confinement reveal fundamental insights into how external mechanical signals influence the organization of the metaphase plate. The flexibility of ECM fibers in our study plays a crucial role in facilitating the natural rounding of mitotic cells by outward deflection of the external fibers, thus promoting the timely progression of mitosis. These observations emphasize the crucial influence of the geometry and flexibility of fibrous ECM environments in shaping mitotic spindle dynamics and ensuring mitotic progression. Our study disambiguates the contributions from RFs and local deformations in the metaphase plate. It shows that the positioning of MP and confinement-driven patterning of RFs on the cortex are critical determinants of mitotic spindle angular positioning within the rounded cell body. Our findings open exciting opportunities to explore confinement-based outcomes in multiple scenarios, including stratification, sprouting, maintenance, growth, and tumoricity^41^. Overall, we identify a new role of fibrous ECM in confinement mitotic biology and shed light on how cells might undergo mitosis with angularly misplaced mitotic spindles *in vivo*.

## Materials and methods

### Generation of nanofiber networks

Aligned and suspended fiber networks were generated from solutions of polystyrene (Scientific Polymer Products, Ontario, NY, USA) dissolved in xylene (X5-500; Thermo Fisher Scientific, Waltham, MA, USA), using our previously reported non-electrospinning Spinneret-based Tunable Engineered Parameters (STEP) technique^55–57^. Force sensing 250 nm fibers were generated from 7wt% solutions of polystyrene (MW: 2,000,000 g/mol) in xylene and deposited with a 12 μm inter-fiber spacing on top of large diameter (2 μm) support fibers prepared from 5 wt% solutions of polystyrene (MW: 15,000,000 g/mol, Agilent Technologies) dissolved in xylene. Fiber networks were bonded at intersection points utilizing solvent vapor in a custom fusing chamber.

### Cell culturing and mitotic synchronization

HeLa cells expressing histone H2B GFP were cultured in Dulbecco’s modified Eagle’s medium (Invitrogen, Carlsbad, CA) supplemented with 10% fetal bovine serum (Gibco, Thermo Fischer Scientific) in T25 flasks (Corning, Corning, NY, USA) and maintained at 37°C and 5% CO_2_ in a humidified incubator. Nanofiber networks were sterilized with 70% ethanol and functionalized with 4 μg/mL of rhodamine-conjugated fibronectin (Cytoskeleton Inc.) in PBS for 1 hr to enable cell-fiber attachment.

Cell division synchronization was performed by treating cells with 9 μM of the Cdk1 inhibitor RO-3306 for 20 h^9^. Cells were subsequently released for the division after 5 times wash with complete culture media.

### Timelapse imaging of mitotic cells

Cells cultured on fibers were imaged every 5 minutes with a 20x 0.8 NA objective in a widefield microscope (Zeiss AxioObserver Z1) equipped with a FITC filter set for GFP fluorescence. Experiments were performed under incubation conditions of 37°C and 5% CO_2_ (Zeiss, Oberkochen, Germany). Mitosis duration was estimated from the timelapse imaging and was taken from the start of the nuclear envelope breakdown (NEBD) to the initiation of telophase.

### Immunofluorescent Staining and Imaging

Cells synchronized with the Cdk1 inhibitor RO-3306 were released with a drug washout and monitored for mitotic entry. Cells were fixed with 4% paraformaldehyde (15 minutes) after ∼ 1 hour after drug washout, where the maximal number of cells were observed (by visual inspection) to be in metaphase. Fixed cells were permeabilized in 0.1% Triton X-100 solution for 15 minutes and blocked with 5% goat serum (Invitrogen, Grand Island, NY) for 45 minutes. Primary antibodies were diluted in an antibody dilution buffer consisting of PBS with 1% Bovine Serum Albumin and Triton-X 100 and stored overnight at 4° C. Primary antibodies include Anti-beta tubulin (1:500, mouse monoclonal, 2 28 33, Invitrogen) and Anti-Hec1 (1:1000, human monoclonal) for labeling microtubules and kinetochores respectively. Secondary antibodies diluted in antibody dilution buffer were added along with the conjugated Phalloidin-TRITC (Santa Cruz Biotechnology) or Alexa Fluor 647 Phalloidin (1:40-1:80, Invitrogen) and stored in a dark place for 45 minutes. Secondary antibodies include donkey anti-human IgG Alexa Fluor 555 (1:600) and Goat anti-mouse IgG Alexa Fluor 405 (1:500, Invitrogen). Confocal microscopy was performed using a laser scanning confocal microscope (LSM 880, Carl Zeiss Inc.) with optimal imaging settings and z-slice thicknesses ranging from 0.36-0.5 μm.

### Analysis of cell parameters

Confocal z-stacks of fluorescently labeled rounded mitotic cells were visualized in the Zeiss Zen software or Image J. The Ortho function in Zen software was utilized to generate the top view (xy) and the cross-sectional front (xz) and side (yz) views. Mitotic cell height and aspect ratio were computed from the cross-sectional yz side views of cells by manual outlining in ImageJ. The cortical perimeter of the cell was estimated from the xz front view of cells.

### Metaphase plate orientation analysis

The metaphase plate (MP) tilt was most evident from the front view of the confocal z-stacks; thus, the xz front views were used to calculate the MP tilt. Measurements were taken with respect to the reference line orthogonal to the plane of the fibers (**Fig. 2a**).

### Retraction fiber analysis

Z-stacks of phalloidin-stained cells were used to quantify the 3D organization of retraction fibers. Mainly, xz front views are generated, and the arc length covered by retraction fibers on either side of the cell (**Fig. 2b**) is measured in ImageJ.

### Quantification of cell forces

Deflections of nanofibers during interphase (contractile cell state, inward fiber deflections) and cell rounding (expansive rounded state, outward fiber deflections) were converted to force (nN) values using our previously reported Nanonet Force Microscopy^29,30,58^. Briefly, fibers are modeled as Euler Bernoulli beams with fixed boundary conditions and subjected to point loads at the cell-fiber contact regions (see *Appendix I* in *Supplementary Materials*).

### Statistical analysis

Statistical analysis was performed in GraphPad Prism (GraphPad Software, La Jolla, CA, USA) software. Statistical comparison among multiple groups was performed using one-way ANOVA and Tukey’s honestly significant difference test. Pairwise statistical comparisons were performed using Student’s t-test. Error bars in scatter data plots indicate standard deviation. *,**,***,**** represent p< 0.05, 0.01, 0.001 and 0.0001 respectively.

### Computational modeling

Simulations are carried out with CSs and CHs represented as particle-like objects positioned at nodes of a cubic lattice confined within a cellular volume. The large lattice structure growing beyond the cell membrane allows the discretization of the cell surface into a finite number of roughly equidistant nodes, similar to the grid size inside the cell. The surface nodes are divided into regions devoid of or coupled to RFs. Using a Monte Carlo simulation, we determine the temporal evolution of CSs and CHs within the cell volume. The potential energies from pairwise interaction forces among CSs, CHs, and the cell membrane are estimated. Achieving the minimum energy ensured a mechanical equilibrium configuration of the spindle. The subsequent equilibrium snapshots are chosen to characterize spindle dynamics and measure chromosomal tilt. For more detailed information on the simulation methods, please refer to *Appendix II* in *Supplementary Materials*.

## ACKNOWLEDGMENTS

ASN acknowledges partial funding support from the National Science Foundation (NSF, Grant No. 2107332 and 2119949). ASN acknowledges the Institute of Critical Technologies and Science (ICTAS) and Macromolecules Innovative Institute (MII) at Virginia Tech for supporting this study. JGD acknowledges funding support from the NIH (R35GM130365). AS and RP acknowledge IACS, Kolkata, for funding support and providing computational facilities.

## Author contributions

ASN conceived the study; AJ, AS, AA, NSG, RP, JGD, and ASN designed research; AJ, JW, and AA performed time-lapse imaging experiments; AJ performed confocal imaging experiments; JW, AJ, AA, ASN, and RK implemented nanonet force microscopy. AS and RP developed computational models and performed simulations; JGD and ASN contributed new reagents/analytic tools; AJ, AA, AS, RP, NSG, JGD, and ASN analyzed data; and AS, AJ, AA, RP, and ASN wrote the paper. All authors contributed to editing and revisions of the paper.

