## Supplementary Materials for "Confinement in fibrous environments positions and orients mitotic spindles"

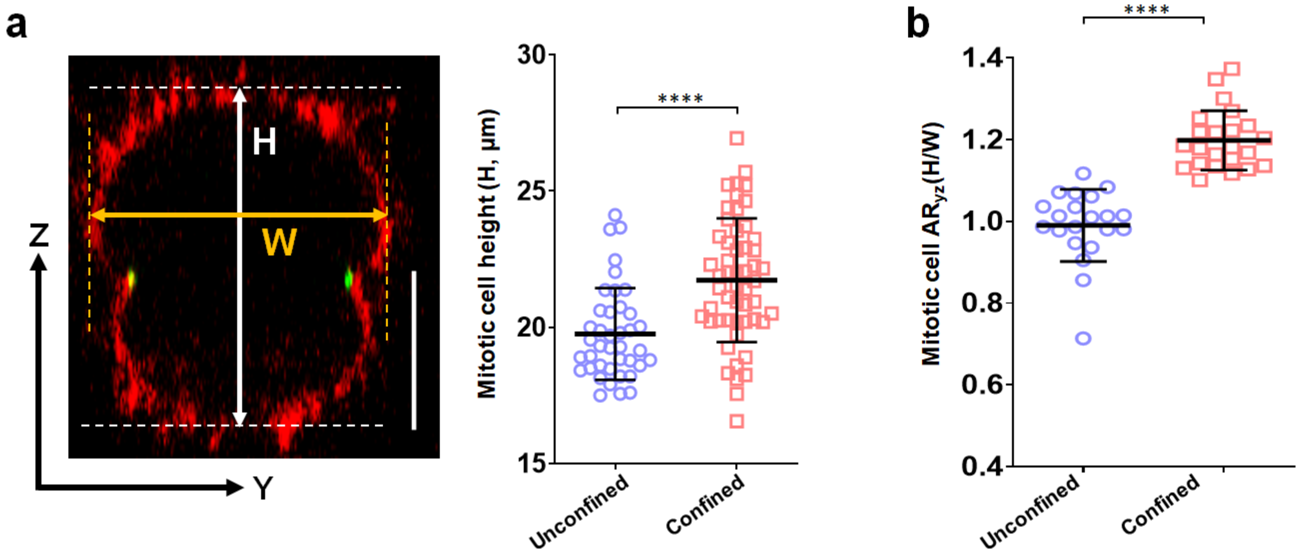


**Fig. S1: Comparison of cell height and aspect ratio in the unconfined and confined categories**. (a) Representative yz cross-section of mitotic cell showing cell height (H) and width (W). Cells in confined categories have more considerable cell height (a, n=40,53 for unconfined and confined respectively) and (b) aspect ratios (b, n=21, 23 for unconfined and confined respectively) than unconfined cases. Scale Bars (10 μm).

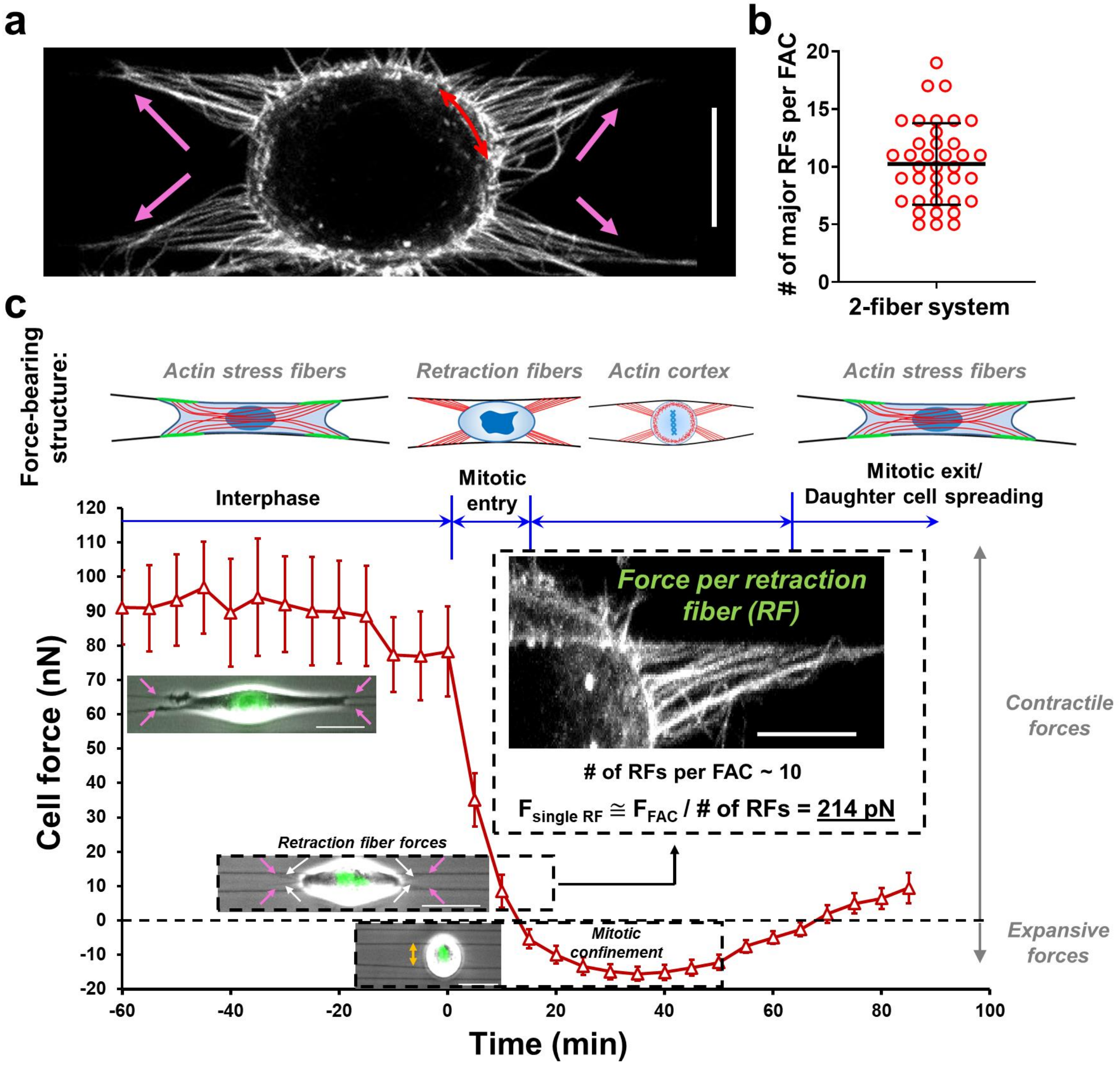


**Fig. S2: Estimation of mechanical tension in individual retraction fibers**: a) Representative stained cell (actin in white) showing 4 sets of retraction fibers, with each set corresponding to a focal adhesion cluster (FAC) during interphase. Scale Bars (10 μm), b) analysis of the number of major retraction fibers associated with each FAC, c) force-bearing structure during different stages before and during mitosis, force per retraction fiber is estimated by taking the force value towards the end of the mitotic entry phase and distributing it equally among the retraction fibers. Scale Bars (10 μm).

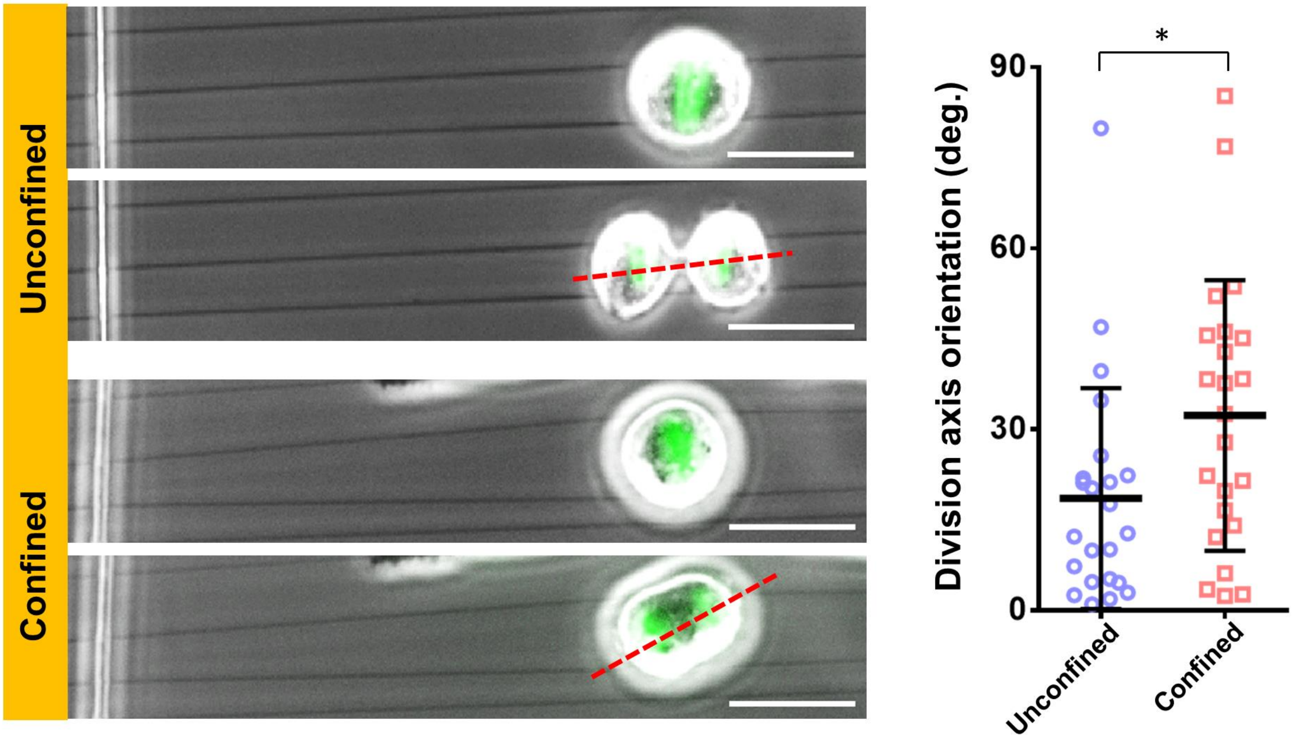


**Fig. S3: Representative images and statistical comparison of division axis orientation for unconfined and confined category cells**; n=23, 24 for unconfined and confined, respectively. Scale Bars (30 μm).

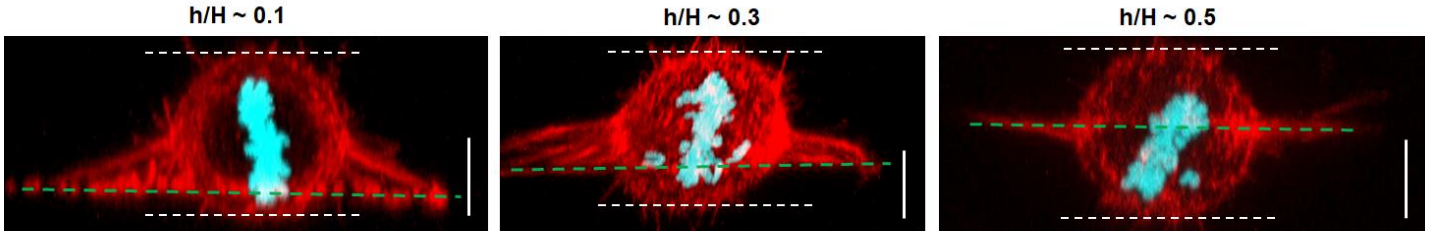


**Fig. S4: Retraction fibers cover the cell cortex area spanning from the fiber plane to the cells' mid-cortical level**; representative front views of cells with h/H ~ 0.1, 0.3, and 0.5, respectively, white dotted lines represent the top and bottom of the cell, the green dotted line represents the external fiber plane. Scale Bars (10 μm).

| Case studied for different RF patterning | Simulated RF pattern | Simulation result | Experimentally consistent/inconsistent |
|---|---|---|---|
| Case 1: RF band displaced with fiber plane | 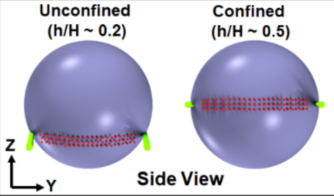  | 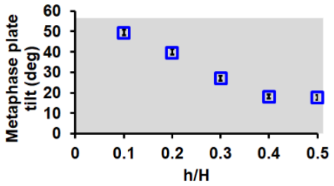  | Inconsistent |
| Case 2: RF band extended up to fiber plane | 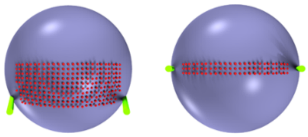 | 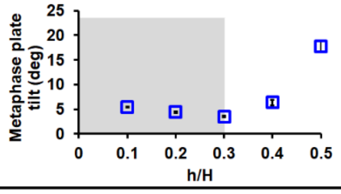  | Inconsistent |
| Case 3: RF band extended as rectangular spots | 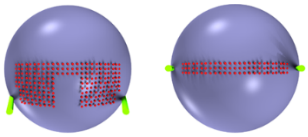 | 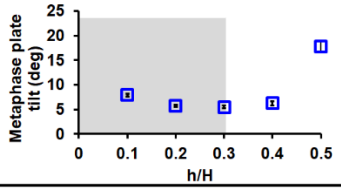  | Inconsistent |
| Case 4: RF-spots (disconnected RF band) extended up to fiber plane | 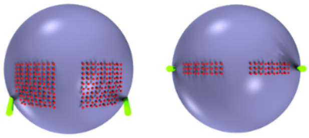 | 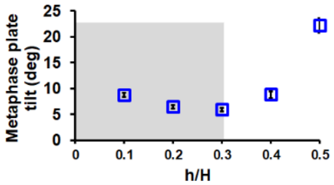  | Inconsistent |
| Case 5: RF band extended as triangles | 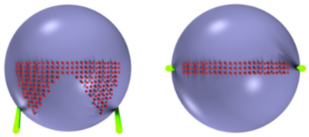 | 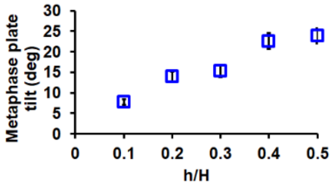  | **Consistent** |
| Case 6: RF spots (disconnected RF band) extended as triangles | 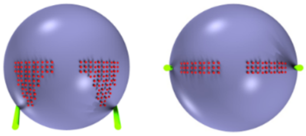 | 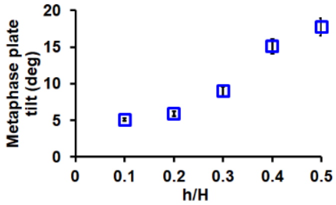  | **Consistent** |

**Table I**: **Different RF configurations produce the experimentally contradictory/consistent MP tilt behavior as a function of confinement (h/H)**. The simulations are performed for the following cases: 1. band-like RF regions that are placed at the fiber plane and displaced with the plane without changing the configurations in both confined and unconfined scenarios, 2. increasing the extent of the RF-band from the mid-cortical level up to the fiber plane, 3. RF-band at the mid-cortical level extended in the form of rectangular spots up to the fiber plane, 4. disjoined spot-like RF arrangements (disconnected RF band) extended from the mid-cortical level as rectangular spots up to the fiber plane, 5. band-like RF arrangements at the mid-cortical level extended as triangular shapes up to the fiber plane, 6. disjoined spot-like RF arrangements (disconnected RF band) extended from the mid-cortical level as triangular shapes up to the fiber plane. The MP tilt in the shaded regions increases with decreasing h/H, contradicting the trend observed in the experiments. Cases 5 and 6 correspond to the experimentally consistent trend of MP tilt angle as a function of mechanical confinement (i.e., h/H) (see Fig. 2 *a, iii* in the main text). The simulation snapshots (column 2) show how the RF organization changes with varying levels of mechanical confinement (h/H), with red dots on the cell surface representing cortical nodes coupled to RFs. Green lines indicate external fibers.

**Different RF configurations produce MP tilt contradictory or consistent with the experimental outcome:**

To investigate if RF arrangements besides triangular patterns in unconfined categories can lead to an experimentally consistent trend of MP tilt with h/H (see Fig. 2 *a*, *iii* in the main text), we ran simulations for a variety of additional configurations (*Table I* in *Supplementary Materials*). When the band of RF moves downward (i.e., decreasing h/H) with the fiber plane without changing its shape and size, the MP tilt angle increases. Due to the downward positioning of the RF band, the resultant pull on the CSs acting from the RFs on the opposite cell surface becomes weaker (*Table I, Case 1* in *Supplementary Materials*). Downward placement of the RF band impedes interaction with the CSs positioned far from the band (often in the cell half not containing the RF bands), resulting in an increased MP tilt. We further find that, by increasing the extent of the equatorial RF-band up to the fiber plane with the downward placement of the fibers, the MP tilt first decreases with decreasing h/H but then increases again below h/H ~0.3, contradicting the experimentally observed behavior (*Table I, Case 2* in *Supplementary Materials*). For h/H greater than 0.3, the effective pull on the CSs adjacent to RF regions on opposite cell surfaces causes a decrease in tilt angle with decreasing h/H. However, when the band region extends at smaller h/H, the CSs in one cell side may stick to the RF regions near the equator. The other CSs on the opposite cell side can localize much below the equatorial plane, causing the MP to be obliquely aligned with the fiber plane. Similar behavior can also be observed with extended rectangular RF spots from the initial band or spot (disconnected band)-like RF arrangements in confined cells with h/H~0.5 (*Table I, Case 3 and Case 4* in *Supplementary Materials*).
Interestingly, the triangular RF regions extended from a band or spot (disconnected band)-like RF arrangements in confined cells can generate experimentally consistent MP tilt (*Table I, Case 5 and Case 6* in *Supplementary Materials*), indicating that the triangular pattern of RFs is critical for experimentally consistent behavior (see Fig. 2 *a, iii* in the main text). Note that, although RF patterning in both Case 5 and 6 (*Table I* in *Supplementary Materials)* can produce experimentally consistent MP tilt, we considered Case 5 to simulate our model. This choice is based on its experimentally observed band-like RF patterning in the confined category (see Fig. 2 *c, i* in the main text).

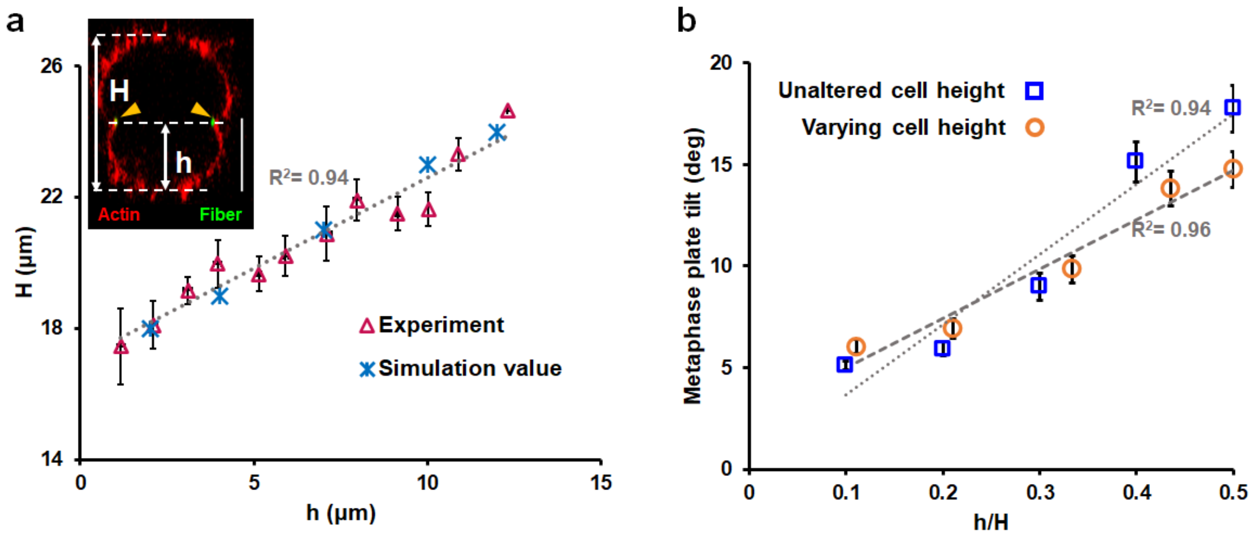


**Fig. S5: The inclusion of experimentally observed cell height variation due to fiber confinement does not significantly alter the behavior of MP tilt vs h/H in simulation.** (a) Blue asterisk points represent the values of h and H seeded in the simulation to find the MP tilt vs. h/H plot in (b) with varying cell height. Errors are the standard error of the mean measured with respect to the corresponding mean values of the data.

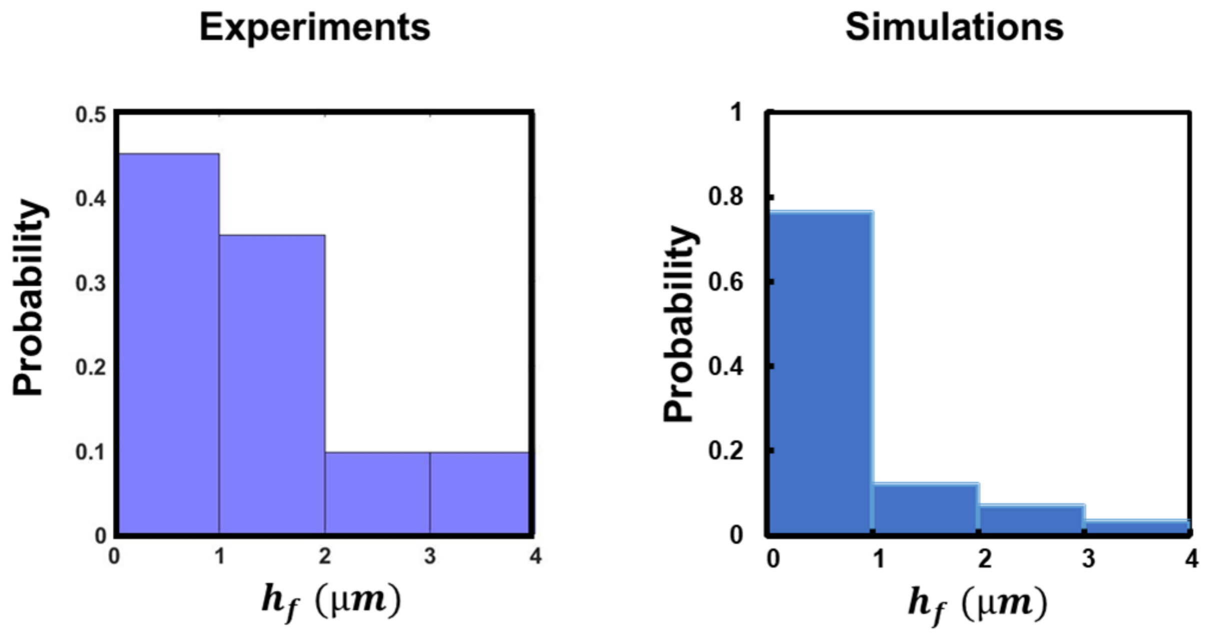


**Fig. S6: Probability distribution of $h_f$ from experimental observations and simulations under high ECM-confinement (h/H ~ 0.5).**

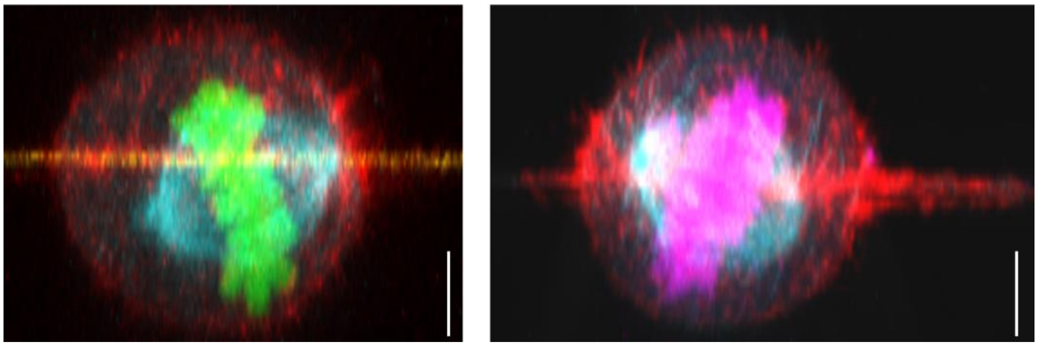

**Fig. S7: Location of spindle poles at high confinement.** Representative front view images showing mitotic spindle pole positioning under high confinement. One pole is near the fiber axis, while the other is rotated and away from the fiber axis, scale bars represent 10 μm.

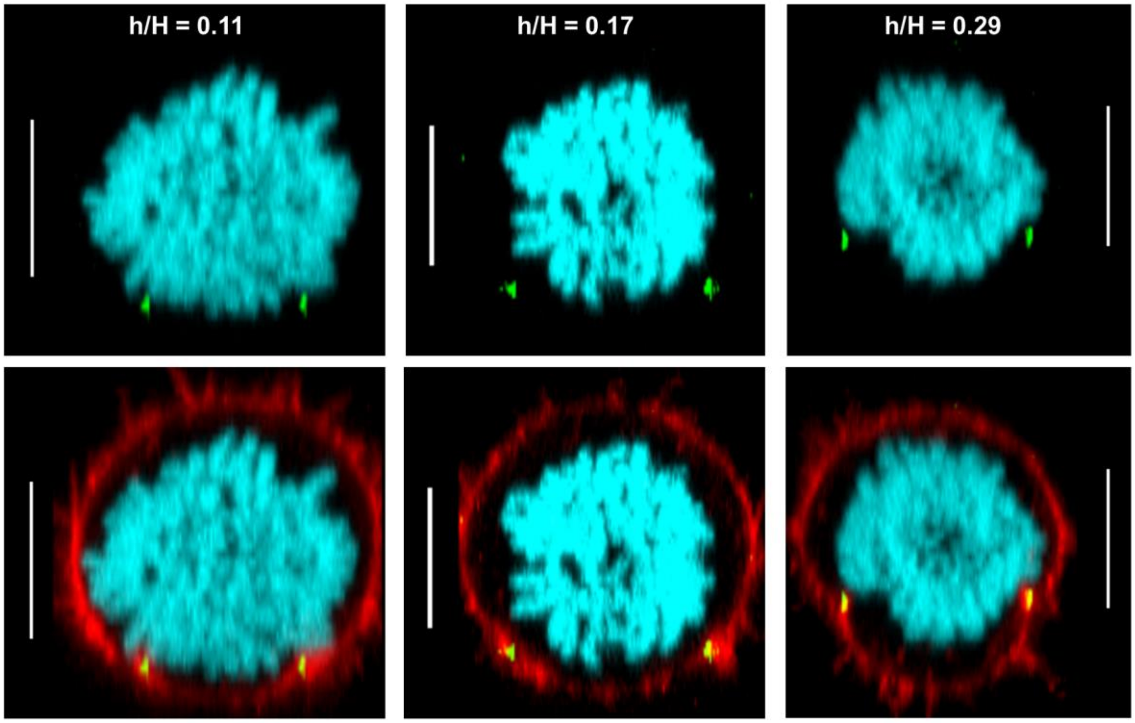


**Fig. S8: Cells at low ECM-confinement do not feature fiber-induced metaphase plate deformations.** Representative yz cross-sections of cells in the unconfined category with different levels of h/H, actin cortex, metaphase plate, and fibers are labeled in red, cyan, and green, respectively; it can be noted that in each case, there is a distinct gap between the spatial location of the metaphase plate and the fibers, scale bars represent 10 μm.

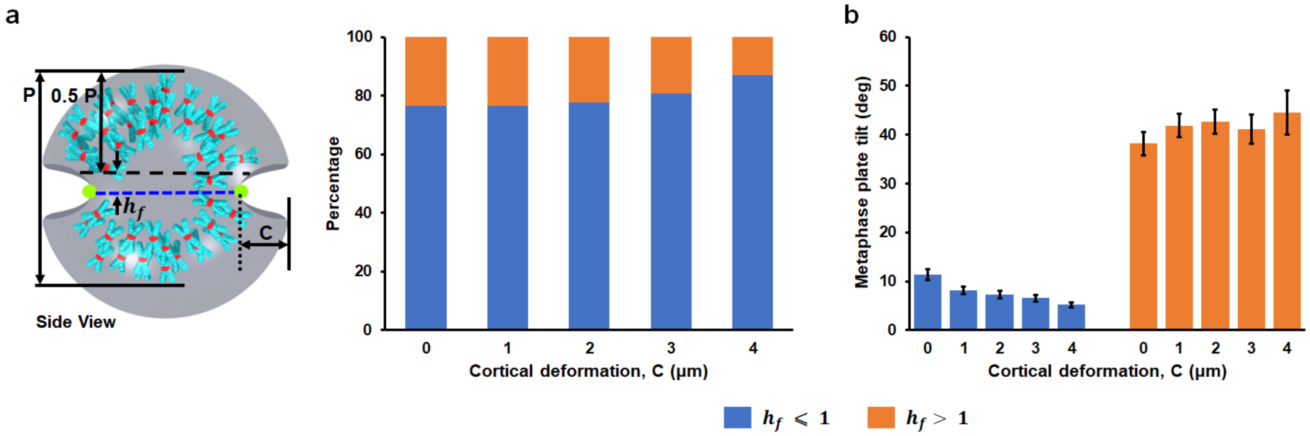


**Fig. S9: Simulation results showing the effect of cortical deformation on MP positioning and its orientation at high ECM-confinement (h/H = 0.5).** (a) Percentage of cells having their MPs positioned symmetrically in both cellular lobes at $h_f \leq 1\ \mu m$, or $h_f > 1\ \mu m$, respectively, for different levels of cortical deformation. Side-view of a representative simulation snapshot in confined cells demonstrating the cortical deformation, C, induced by the external fibers and $h_f$ metric, denoting the distance between the fiber plane and the MP center. 'P' is the length from the side view of MP, and 0.5 P defines the MP mid-plane. CHs are blue, KTs are red, and two green dots denote external fibers. (b) Metaphase plate tilt vs. cortical deformation for the MP positioned at $h_f \leq 1\ \mu m$, or $h_f > 1\ \mu m$, respectively. MP tilt for $h_f \leq 1\ \mu m$ decreases with higher levels of cortical deformation. However, there is no clear trend observed for $h_f > 1\ \mu m$.

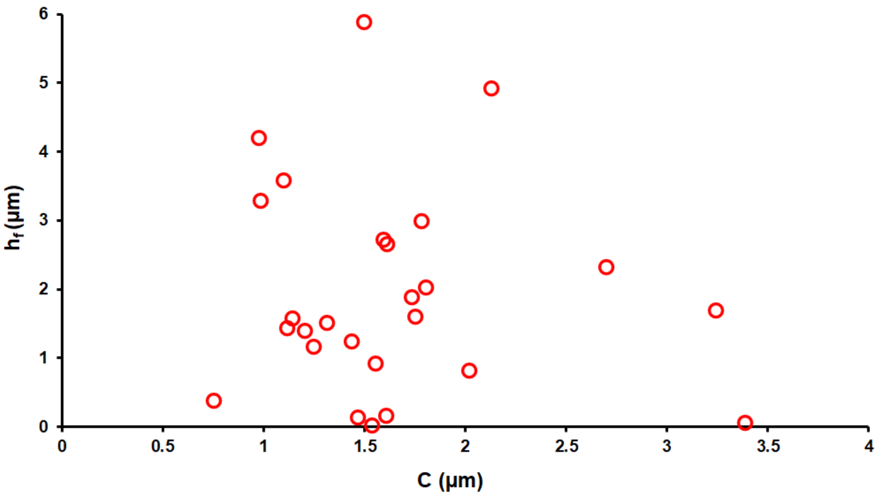


**Fig. S10: Experimental data demonstrating a weak correlation between cortical deformation (C) and positioning of the MP with respect to the fiber plane ($h_f$).**

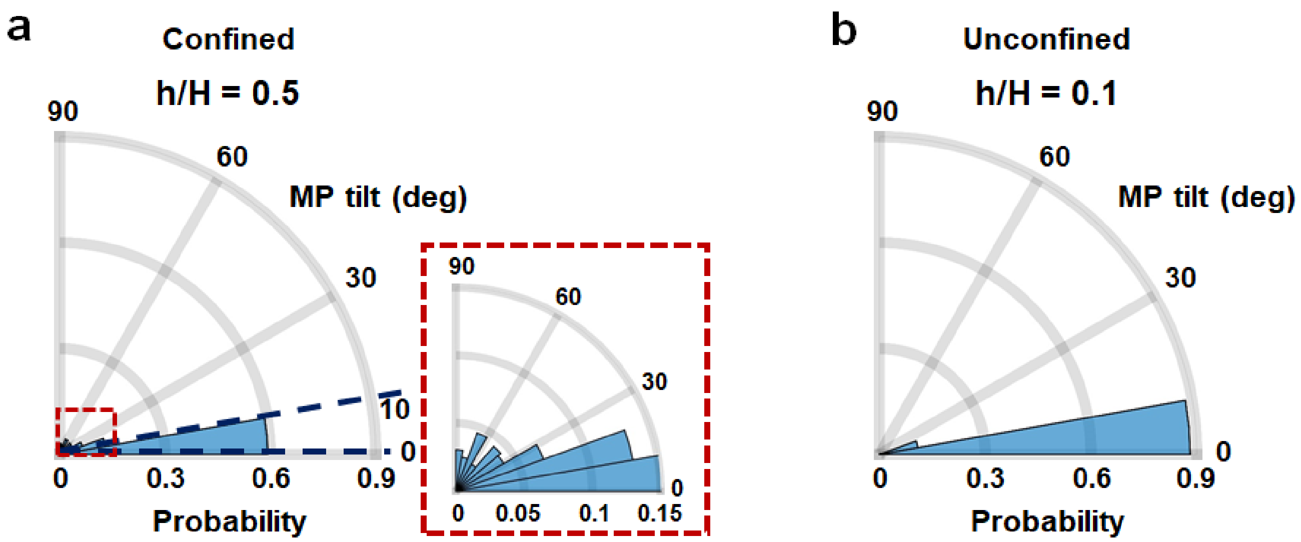


**Fig. S11: Polar histogram of MP tilt angle in spherical confined and unconfined cells.** Inset in *a* shows a magnified view of the polar histograms, demonstrating a comparatively larger fluctuation in MP tilt angle in confined cells than in unconfined conditions.

**Supplementary Movie S1:** 3-dimensional rendering of a confocal z-stack showing a rounded HeLa cell in metaphase, positioned on top of a fiber doublet.

**Supplementary Movie S2:** 3-dimensional rendering of a confocal z-stack showing a rounded HeLa cell in metaphase, mechanically confined within a fiber doublet. Actin: red, Fibers: green and chromosomes: blue.

**Supplementary Movie S3:** HeLa cell expressing Histone H2B GFP, entrapped in fiber doublets deflecting fibers outward during mitotic rounding (timestamp-h:min) and scale bar 20 µm.

**Supplementary Movie S4:** HeLa cell dividing under confinement, showing tilted orientation of division axis (timestamp-h:min) and scale bar 20 µm.

### APPENDIX

#### I. NANONET FORCE MICROSCOPY

We used our previously reported 'Nanonet force microscopy' to quantify cell forces at various cell cycle stages. We employ nanofiber networks consisting of large diameter (2 µm) support fibers placed ~350 µm apart and an orthogonal layer of small diameter (250 nm) fiber layer with an inter-fiber spacing of ~ 12 µm. The fiber layers are fused at their junctions through solvent vapor exposure. This leads to the generation of fixed-fixed boundary conditions at both ends of the small-diameter fiber. Fibers are modeled as Euler-Bernoulli beams. Cellular force exertion on fibers depends on the cell cycle stage.

Since focal adhesion organization demonstrates distinct clustering at the cell extremities during interphase, the force exertion on each fiber can be considered a 2-point load. In this stage, cells exert horizontal and vertical forces on the fibers. The direction of this resultant force exertion is taken along the average orientation of the actin stress fibers ($\alpha_{SF}$) emerging from each focal adhesion cluster. During mitotic entry, the breakdown of actin stress fibers occurs, and contractile forces (inward fiber deflection) are generated by the actin-based retraction fibers and their orientation ($\alpha_{RF}$) determines the direction of resultant force exertion. The actin stress and retraction fibers' orientation are quantified from the immunofluorescent staining of actin.

During mitosis, cells adopt a rounded shape and push fibers outward. In this configuration, the force exertion can be estimated by using a vertical force model, with the expansive forces from the stiff actin cortex acting orthogonal ($\alpha = 90^0$) to the fibers.

| Stage in cell cycle | Cytoskeletal/ adhesion organization | Force-bearing structure | Direction of cell forces |
|---|---|---|---|
| *Interphase* | 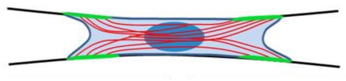 | *Actin stress fibers* | 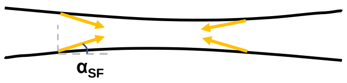  |
| *Mitotic entry* | 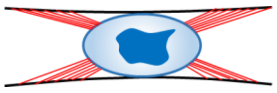 | *Actin-based retraction fibers* | 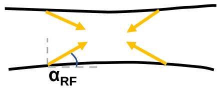  |
| *Mitosis* | 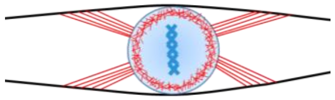 | *Actin cortex* | 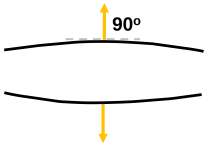  |

Characterization of forces at different stages of the cell-cycle based on cytoskeletal organization

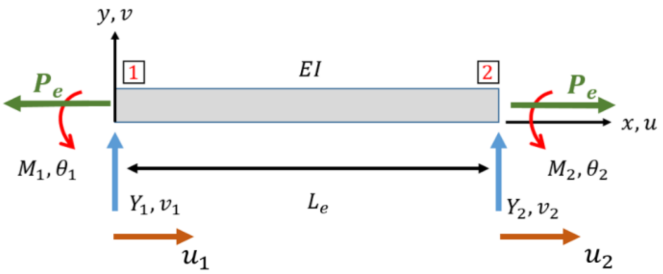


Finite beam element with axial force ($P_e$), shear force (Y) and bending moment (M)

For force quantification, we quantify the cell-mediated fiber deflections, and these deflection profiles of fibers bonded to the support fibers on either end serve as input for the optimization framework to calculate the cell forces. Specifically, the '*load updating for finite element models*' method has been utilized, and the optimization to determine cell forces is performed through a MATLAB gradient-based optimizer (GBO).

The formulation for the cell forces has been reported previously by our group [1, 2]. Each taut fiber can be estimated as a beam with an axial force. The Finite Element Method (FEM) can be used to discretize the beam into a finite number of uniform straight beam elements, as shown in this schematic. Shear forces and moments at two nodes are indicated by $Y_1, M_1, Y_2$ and $M_2$.

Using Castigliano's theorem to model the strain energy of each beam element, we can derive the interrelationship between the shear forces and moments exerted on the beam elements and the corresponding displacements.

$$\{\boldsymbol{F}\} = \left[[\boldsymbol{k}] + [\boldsymbol{n}]\right]\{\boldsymbol{q}\}$$

Here, $[\boldsymbol{k}]$ and $[\boldsymbol{n}]$ will be referred to as the fundamental stiffness matrix and incremental stiffness matrix or geometric stiffness matrix, respectively.

Here,

$$\{\boldsymbol{F}\} = \begin{Bmatrix} Y_1 \\ M_1 \\ Y_2 \\ M_2 \end{Bmatrix}, [\boldsymbol{k}] = \frac{EI}{Le}\begin{bmatrix} \frac{12}{Le^2} & \frac{6}{Le} & -\frac{12}{Le^2} & \frac{6}{Le} \\ \frac{6}{Le} & 4 & -\frac{6}{Le} & 2 \\ -\frac{12}{Le^2} & -\frac{6}{Le} & \frac{12}{Le^2} & -\frac{6}{Le} \\ \frac{6}{Le} & 2 & -\frac{6}{Le} & 4 \end{bmatrix},$$

$$[\boldsymbol{n}] = \frac{P_e}{10}\begin{bmatrix} \frac{12}{Le} & 1 & -\frac{12}{Le} & 1 \\ 1 & \frac{4Le}{3} & -1 & -\frac{Le}{3} \\ -\frac{12}{Le} & -1 & \frac{12}{Le} & -1 \\ 1 & -\frac{Le}{3} & -1 & \frac{4Le}{3} \end{bmatrix}, \{\boldsymbol{q}\} = \begin{Bmatrix} v_1 \\ \theta_1 \\ v_2 \\ \theta_2 \end{Bmatrix}$$

$P_e$ is the axial tensile force in the element and $L_e$ is the length of the element.

The optimization framework is based on minimizing the following objective function.

$$g(x) = \frac{1}{2}\left|\left|\boldsymbol{V}_{\boldsymbol{EXP}} - \boldsymbol{V}_{\boldsymbol{FEM}}\right|\right|^2$$

Here, $\boldsymbol{V}_{\boldsymbol{EXP}}$ represents the vector of vertical displacements generated from interpolation of the experimental vertical displacements and $\boldsymbol{V}_{\boldsymbol{FEM}}$ is the vector of computational vertical displacements from the finite element model.

The overall flowchart for the optimization routine is as follows:

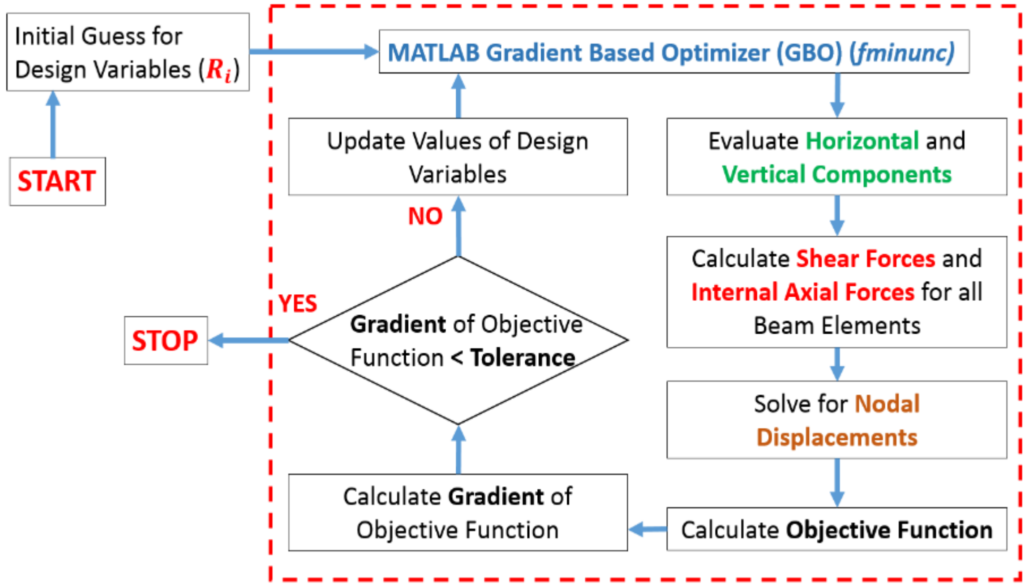


The tolerance is set at $10^{-10}$ and once the optimization criteria are reached, the cell forces corresponding to a particular fiber deflection profile are characterized.

The following properties of the beam are used for cell force calculations [2, 3].

| Property | Value |
|---|---|
| Length (µm) | 350 |
| Diameter (nm) | 250 |
| Young's Modulus (GPa) | 0.97 |
| Pre-Tension (nN) | 201.2 |

#### II. CELL COMPUTATIONAL MODEL

In our model, centrosomes (CSs) and chromosomes (CHs) are distributed randomly within a spherical cell of radius $R_c$. The cell's interior is divided into a three-dimensional cubic lattice grid with a lattice spacing of 0.5 $\mu m$. The interaction of the lattice structure with the cell influences the discretization of the cell surface, resulting in roughly equidistant nodes with inter-node distances comparable to the lattice spacing. The cell membrane, consisting of discrete nodes, is divided into two distinct regions. One part consists of areas where retraction fibers (RF) attach to the cell surface, connecting it to the external parallel fibers forming the cortical RF zones. On the other hand, the remaining regions are devoid of such RF attachments. Note that our model does not account for the elasticity of the cell membrane and the stiffness of the actin cortex.

##### A. Construction of RF-regions and the cortical deformation

$R_c$, θ, and φ parameterize the spherical cell. $\theta$ (∈ 0,180$^0$) represents the polar angle, measured with respect to the positive *z*-axis, and $\varphi$ (∈ 0,360$^0$) represents the azimuthal angle, measured from the positive *x*-axis. In our model, depending upon the cellular position with respect to external fibers, we have incorporated the shapes of the RF regions on the cell surface based on experimental findings in Fig. 2 *c, i*. For the purely confined scenario ($h/H = 0.5$), we considered two bands of RF regions, each appearing on opposite cell surfaces. The center of the bands is located at $\varphi_0 = 0$ (along the positive *x*-axis) and 180$^0$ (along the negative *x*-axis), respectively, while placed at the cell equatorial plane at $\theta = 90^0$. The bands' width and extent along the cell surface are governed by $d\theta$ and $d\varphi$, respectively. Therefore, the bands are ranges from $\theta \in 90^0 - d\theta/2,\ 90^0 + d\theta/2$ and $\varphi \in \varphi_0 - d\varphi/2,\ \varphi_0 + d\varphi/2$, respectively. As we move towards the unconfined circumstances, the bands are extended in the form of inverted triangles, with the vertex of the triangles moving downward and reaching the fiber plane (see simulated RF pattern in *Table I, Case 5* in *Supplementary Materials*). Furthermore, cortical deformation is introduced in the model by placing two thin cylinders at opposite cell edges, with the axes of the cylinders lying on the external fiber plane (see simulation snapshot in *Supplementary Fig. S9 a*).

##### B. The interaction forces and the corresponding potential energies

We treat the CSs and CHs as particle-like objects interacting with each other and the cell surface nodes through distance-dependent forces [4-8]. However, the interaction between CSs and the kinetochores (KTs) located at the centromeric regions of CHs is assumed to be distance-independent [4, 5, 9].

Organelles interact via dynamic microtubules. The forces between organelles are considered to decay exponentially with distance due to a falling microtubule density. This assumption is based on the simplified notion that microtubule length distribution follows an exponential pattern, resulting in a spatially decaying exponential force profile [7, 10, 11]. The forces under consideration are conservative; thus, interaction among each pair of objects corresponds to a potential energy. The forces between a pair of objects (CS, CH/KT) as a function of distance $r$ and the corresponding change in potential energy of shifting one object from another changing the distance between them from $r_1$ to $r_2$ can be computed as follows:

$$f_{CS-CS} = f_{CS-CS}^{(0)} e^{-r/L_1}, \qquad \Delta E_{CS-CS} = -f_{CS-CS}^{(0)} \int_{r_1}^{r_2} e^{-r/L_1}\, dr. \tag{1}$$

$$f_{CS-KT} = f_{CS-KT}^{(0)}, \qquad \Delta E_{CS-KT} = -f_{CS-KT}^{(0)} \int_{r_1}^{r_2} dr\ . \tag{2}$$

$$f_{CS-CH} = f_{CS-CH}^{(0)} e^{-r/L_1}, \qquad \Delta E_{CS-CH} = -f_{CS-CH}^{(0)} \int_{r_1}^{r_2} e^{-r/L_1}\, dr. \tag{3}$$

Here, $f_{CS-CS}^{(0)}, f_{CS-KT}^{(0)}$, and $f_{CS-CH}^{(0)}$ represent the amplitude of the forces between each CS-CS, CS-KT, and CS-CH pairs, respectively, and $L_1$ is the spatial range of respective forces.

Similarly, the forces acting on individual CSs from each surface node point with coordinate $\mathbf{s}$, as well as the corresponding potential energy of moving a CS from a distance $r_1(\mathbf{s})$ to $r_2(\mathbf{s})$ away from the surface nodes, can be computed to find the overall potential energy change due to the regions of the cell surface lacking the RF attachments or coupled to RFs, as follows:

$$f_{CS-CRTX} = f_{CS-CRTX}^{(0)} e^{-r(\mathbf{s})/L_2}, \qquad \Delta E_{CS-CRTX} = -\int_{\Omega} d\mathbf{s} \int_{r_1(\mathbf{s})}^{r_2(\mathbf{s})} f_{CS-CRTX}^{(0)} e^{-r/L_2}\, dr. \tag{4}$$

$$f_{CS-RF} = f_{CS-RF}^{(0)} e^{-r(s)/L_2}, \qquad \Delta E_{CS-RF} = -\int_{\Omega'} d\mathbf{s} \int_{r_1(\mathbf{s})}^{r_2(\mathbf{s})} f_{CS-RF}^{(0)} e^{-r/L_2}\, dr. \tag{5}$$

Here, the amplitudes of the forces from cortical nodes devoid of or coupled to RFs are denoted by $f_{CS-CRTX}^{(0)}$, and $f_{CS-RF}^{(0)}$, respectively. The interaction between CSs and cell membranes is assumed to be short-ranged, resulting in CSs interacting strongly with the adjacent cell surface regions, with $L_2$ representing the spatial ranges of the respective force interactions [4, 5]. The integration over the cortical region devoid of RFs (Ω) and attached to RFs (Ω') can be expressed as a sum over discrete surface nodes.

To maintain the majority of the spindles bipolar, each CS exerts attractive forces on other CSs, KTs, and the cell membranes while repelling the CH arms [4, 5]. These forces arise from the interaction of CS-nucleated MTs with another CS, possibly via Dynein or Kiensin-14, MTs connecting CS to KTs and cell membranes via Dynein or pushing the chromosomal arms via chromokinesins (such as kinesin-10 and kinesin-4) [7, 12-17]. The amplitudes of the forces are assumed to be proportional to the average number of motors operating at the MTs-organelle interface.

The interaction between pairs of chromosomes involves a steric repulsion that scales as the inverse square of their mutual distance. The repulsion force between pairs of chromosomes becomes effective when their mutual distance falls below a specific threshold set to be less than 2 $\mu m$ in the simulation. Also, note that there are no explicit chromosomal arms in the simulation. The arms shown in the simulation snapshots are for visualization purposes only. We use two force vectors for a single chromosome: one linking the CS and KT and the other parallel vector connecting the CS to the corresponding chromosome arms. We use the centers of mass of both KTs and chromosomal arms to be positioned around the centromeric region of the chromosome.

##### C. Monte-Carlo Simulation

The Monte Carlo algorithm is employed to simulate the temporal evolution of the model components, governed by the above-mentioned force interactions. Initially, 8 CSs and 46 CHs are randomly placed on the lattice grid points within the cell. The CSs and CHs are constrained to move within the cell volume. CS, or CH, cannot occupy the position that is already occupied by CSs/CHs.

During each Monte Carlo step, the system undergoes the following updates:

1. A CS or CH is selected randomly.
2. If the selected object is a CS, the net energy change ($\Delta E$) associated with moving the CS from its current position, $r_1$, to a randomly selected unoccupied neighboring position, $r_2$, is computed by adding $\Delta E_{CS-CS}$, $\Delta E_{CS-KT}$, $\Delta E_{CS-CH}$, $\Delta E_{CS-CRTX}$, and $\Delta E_{CS-RF}$, respectively.
3. If the selected object is a CH, the energy change ($\Delta E$) is determined by considering only the contributions from $\Delta E_{CS-KT}$ and $\Delta E_{CS-CH}$.

The move is accepted if $\Delta E \leq 0$. If $\Delta E > 0$, the move can still occur based on the Boltzmann weight, calculated as $P = e^{-\beta\Delta E}$, where $\beta$ is inversely proportional to the effective temperature required for updating the system configuration [18]. The simulation continues until a stable mechanical equilibrium is achieved. In the equilibrium configuration, we consider two or more CSs clustered if found in the neighboring lattice points. We simulate $500-1000$ different random initializations of the model configurations. The specific model parameters can be found in Table II, and more detailed information regarding the simulation methodology can be found in [4] and [5].

**Table II. List of parameters**

| Abbreviations | Meaning | Value |
|---|---|---|
| $\boldsymbol{f_{CS-CS}^{(0)}}$ | Amplitude of CS-CS attraction | $-1.7\ pN$ |
| $\boldsymbol{f_{CS-KT}^{(0)}}$ | Amplitude of CS-KT attraction | $-4.0\ pN$ |
| $\boldsymbol{f_{CS-CH}^{(0)}}$ | Amplitude of CS-CH repulsion | $10\ pN$ |
| $\boldsymbol{f_{CS-CRTX}^{(0)}}$ | Amplitude of CS-CRTX attraction | $0.09\ pN$ |
| $\boldsymbol{f_{CS-RF}^{(0)}}$ | Amplitude of CS-RF attraction | $7 \times\ f_{CS-CRTX}^{(0)}$ |
| $\boldsymbol{R_c}$ | Cell radius | $10\ \mu m$ |
| $\boldsymbol{d\theta, d\varphi}$ | Annular widths of the RF bands in confined cells | $8^0, 75^0$ |
| $\boldsymbol{L_1}$ | Spatial range of CS-CS and CS-CH forces | $1.25\ \times\ R_c$ |
| $\boldsymbol{L_2}$ | Spatial range of CS-CRTX and CS-RF forces | $\frac{L_1}{4}\mu m$ |
| $\boldsymbol{\beta}$ | Inverse temperature | $20\ (pN\ \times \mu m)^{-1}$ |
